# How exposure to land use impacts and climate change may prune the tetrapod tree of life

**DOI:** 10.1101/2022.02.01.478740

**Authors:** Linda J Beaumont, David A Nipperess, Peter D Wilson, John B Baumgartner, Manuel Esperon-Rodriguez

## Abstract

Human domination of landscapes is a key driver of biodiversity loss, with the fingerprint of climate change becoming increasingly pronounced. Frameworks and tools for identifying threats to biodiversity are required to meet Post-2020 Global Biodiversity Framework targets for 2030 that call for, among other things, reducing or halting species extinction rates (*1*). Hence, we compiled a phylogenetic tree for terrestrial tetrapods, mapped hotspots of geographically restricted and evolutionarily distinct lineages, and identified which hotspots may simultaneously face the highest magnitudes of land use impacts and climate change. Across a quarter of Earth’s surface, hotspots contain the entire ranges of 45% of tetrapods, representing 39% of terrestrial tetrapod evolutionary heritage. By 2070, we estimate 8–13% of this heritage to occur entirely within hotspots highly exposed to climate change, with 13–29% of hotspots projected to experience high exposure to both stressors simultaneously. Most hotspots at highest risk occur in countries least able to take action. Our analysis highlights the need for global ambition and coordination to avoid catastrophic loss of tetrapod evolutionary heritage.

## Main Text

Over the 35+ years since Soulé described conservation biology as a “mission-oriented crisis discipline” (*2*), and despite global initiatives including the Millennium Ecosystem Assessment (*3*) and the Convention on Biological Diversity’s Aichi Biodiversity Targets (*4*), the state of the world’s biodiversity continues to decline (*5*). Approximately one quarter of species assessed by the IUCN are threatened with extinction and population sizes of vertebrates have declined on average 68% over 1970–2016 (*6*), with extinctions exceeding the background rate by orders of magnitude (*7, 8*). Terrestrial biodiversity decline has predominantly been driven by over-exploitation, habitat loss, fragmentation and degradation (*9*), with one-third of the world’s terrestrial surface utilized for agriculture, and 75% of ice-free land having undergone significant human-caused transformation (*6*).

In coming decades, climate change will reconfigure and create new climate zones, resulting in spatial shifts to the bioclimatic envelopes that ultimately govern species’ distributions (*10*). Multiple lines of evidence portend widespread rearrangement of terrestrial, freshwater and marine communities by the end of this century (*11, 12*) which, in conjunction with species extinctions (*13, 14*), will impact the functioning of natural ecosystems and the well-being of human populations (*15, 16*).

Prioritizing actions to reduce threats and safeguard biodiversity requires that we identify regions rich with biological heritage and assess the extent to which these regions are under pressure from anthropogenic stressors (*17*). Here, we identified and mapped the most biologically irreplaceable regions, which we define as hotspots of geographically restricted and evolutionarily distinct terrestrial tetrapod lineages. We then assessed risk to the biodiversity within these regions caused by exposure to land use impacts and/or climate change. Risk was determined by the extent to which regions are projected to be exposed to these stressors individually and combined, for the periods 2050 and 2070. With this approach, we have identified regions where conservation intervention may have the greatest impact on the tetrapod tree of life.

### Hotspots of geographically restricted and evolutionarily distinct tetrapod lineages

What makes a location important to conservation is its relative contribution to global evolutionary heritage (*18*). As such, the more evolutionary lineages uniquely or mostly represented at a site, the more irreplaceable the site is (*19*). To measure irreplaceability, we compiled the most complete terrestrial tetrapod phylogenetic tree to date (n = 33,199 species) by combining separate trees for mammals (Mammalia), birds (Aves), squamate reptiles (Squamata), crocodiles (Crocodylia), turtles (Testudines), tuatara (Rhynchocephalia) and amphibians (Lissamphibia) (**Fig. 1**). Using expert range maps (*20*–*22*), we assessed the incidence of each tetrapod species in each 10 km x 10 km grid cell (n = 1,340,520) across the Earth’s land surface, then summed the corresponding branch lengths of the phylogenetic tree that are unique to a given cell when compared to 1000 randomly selected cells (i.e., complementarity in Phylogenetic Diversity (*23*)). We refer to regions with the highest 25% of complementarity values as *Very Important Places* (VIPs) (n = 335,130 grid cells) (**Fig. 2**). (Different thresholds in complementarity can be explored through our VIP-Explorer ShinyApp, **data S1**).

**Fig. 1.**
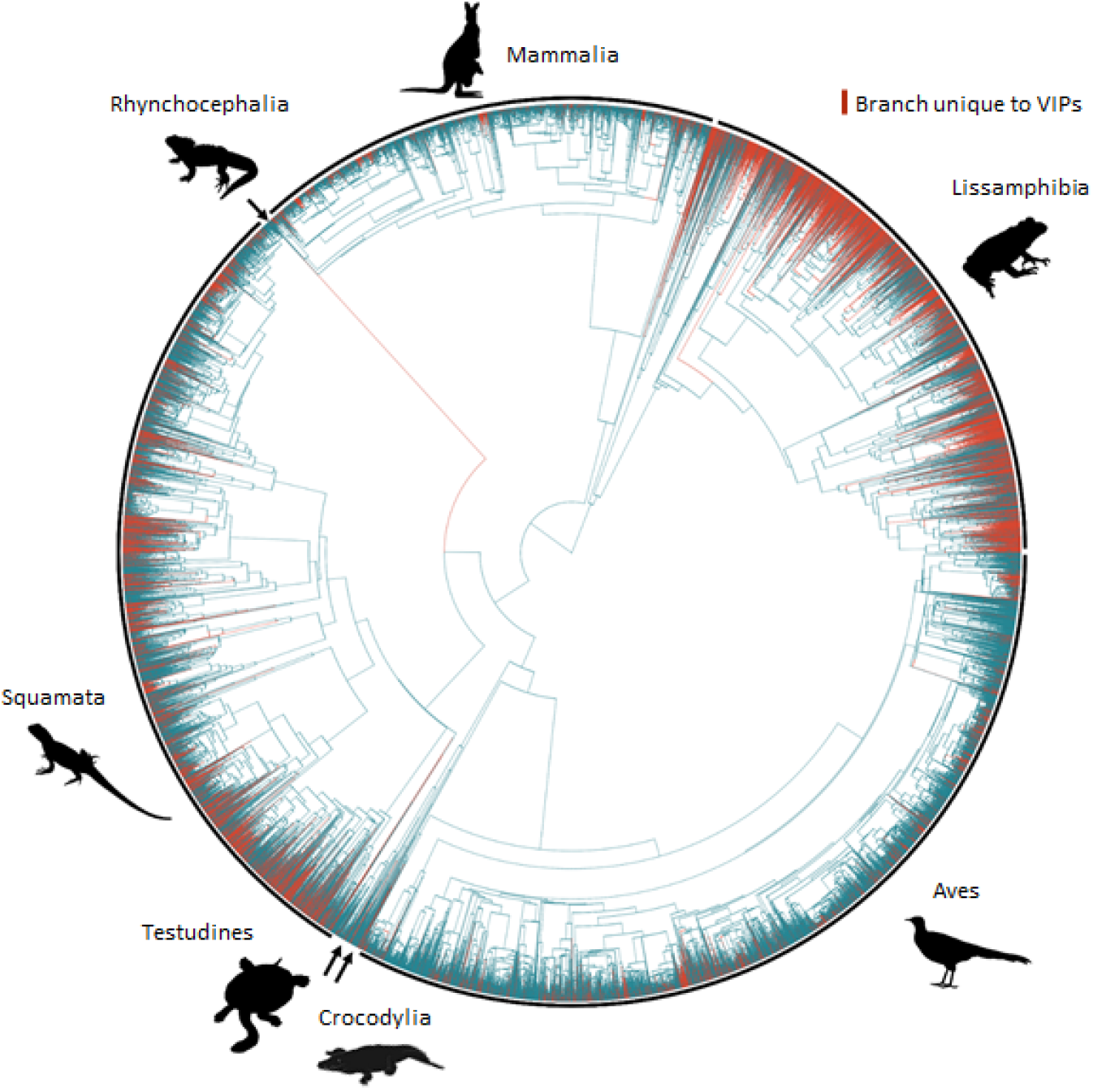
Phylogenetic tree of the world’s terrestrial tetrapod species. Phylogenetic tree of 33,199 tetrapod species (mammals [Mammalia], birds [Aves], squamate reptiles [Squamata], crocodiles [Crocodylia], turtles [Testudines], tuatara [Rhynchocephalia] and amphibians [Lissamphibia]), with highlighted branches being restricted to *Very Important Places* (VIPs) (i.e., 25% of the terrestrial surface with the highest concentration of geographically restricted and evolutionarily distinct tetrapod lineages). Silhouettes sourced from phylopic (www.phylopic.org).

**Fig. 2.**
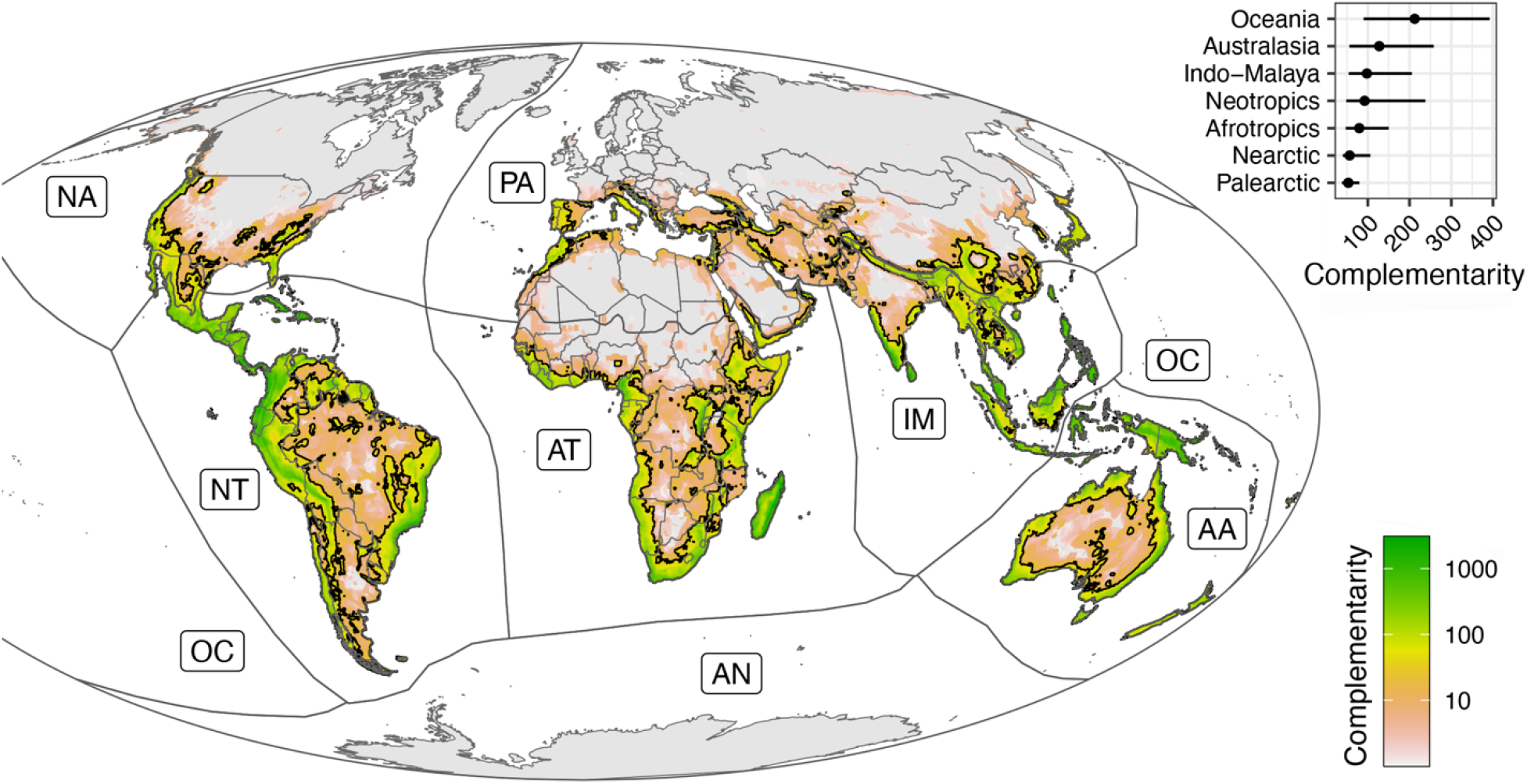
Hotspots of geographically restricted and evolutionarily distinct tetrapod lineages. Map (Mollweide projection) showing the complementarity in tetrapod Phylogenetic Diversity of each 10 km cell, when compared to 1000 randomly selected cells. The colour scale represents Phylogenetic Diversity (i.e., Complementarity) in millions of years. The outline represents the quarter of the earth’s terrestrial area with the highest complementarity, i.e., hotspots of geographically restricted and evolutionarily distinct tetrapod lineages. We refer to these regions as *Very Important Places* (VIPs). Inset: Median and Interquartile range of complementarity values for grid cells in each biogeographic realm. AA = Australasia; AT = Afrotropics; IM= Indo-Malaya; NA= Neartic; NT= Neotropics; PA = Paleartic; OC = Oceania.

We found that almost half of the world’s terrestrial tetrapod species (45%), representing 39% of terrestrial tetrapod evolutionary heritage, are restricted to VIPs, with squamate reptiles and amphibians making larger contributions to VIPs than mammals and birds (**Fig. 1)**. Of the world’s seven biogeographic realms, VIPs in Oceania have the highest median irreplaceability (212 IQR: 302) followed by Australasia (127 IQR: 202), with the Palearctic having the lowest (53 IQR: 42) (**Fig. 2**). The highest number of VIP grid cells, however, occur in the Tropical/Subtropical Moist Broadleaf Forests (38%) of the Neotropics, Indo-Malaya and Afrotropics, as well as the Tropical and Subtropical Grasslands, Savannas and Shrublands (15%) mostly in the Afrotropics, Australasia and Neotropics (**table S1**). Sizeable tracts of VIPs also occur within the Deserts and Xeric Shrublands (13%), and the Temperate Broadleaf and Mixed Forests (8%) of the Palearctic and Australasia.

The regions we identified as VIPs have been previously highlighted for their rich biodiversity (*24, 25*) and importance for restoration and/or conservation (*24, 26*). VIPs occur in higher concentration at lower latitudes, especially along steep environmental gradients (such as in mountainous terrain) and/or geographically isolated landmasses. These factors are known to correlate with species having small ranges (*27*), contributing to the relative irreplaceability of these locations. Further, the smaller ranges of squamate reptiles and amphibians in comparison to other clades (**fig. S1**), combined with phylogenetic clustering of small ranges and longer terminal branches, means that the presence of these taxa increases the irreplaceability value of a grid cell.

The accumulation of distinctive evolutionary lineages in VIPs can also be attributed to both geographic isolation, contributing to faunal provincialism (*28*), and long periods of relative environmental stability resulting in ‘museums’ of persistent lineages (*29*), especially in the tropics (*30*). The Southern Hemisphere is disproportionately represented among VIPs because of a long history of continental isolation, engendering significant indigenous radiations of tetrapod lineages (*31*). There is also variation in the location of VIPs for different clades, reflecting, in part, differing capacities to disperse between isolated landmasses (**data S1**).

### A global assessment of risks to VIPs from exposure to land use impacts and climate change

We assessed loss of habitat condition due to land use impacts predicted for 2050 and 2070, by adapting data from the Land Use Harmonization (LUH2) datasets that were developed as part of the Land-Use Model Intercomparison Project (*32*). We evaluated projections under two Shared Socioeconomic Pathways, SSP2 and SSP5, which connect to two Representative Concentration Pathways (RCP), RCP 4.5 and RCP 8.5, respectively. Each of the land-use categories in LUH2, expressed as the proportion of land in a 0.25 × 0.25 degree cell, was scaled according to habitat quality (*33*) (**table S2**) and summed to produce a habitat condition score. The scaling coefficients estimate the proportion of plant species richness remaining for that land use class after conversion from a pristine state (*33*). The habitat condition score was subtracted from 1, to represent potential biodiversity loss due to land use impacts.

We estimated exposure to climate change by measuring the difference between average baseline multidimensional climate (1979–2013) in grid cells and future climate predicted by an ensemble of global climate models. We obtained climate data from the CHELSA data set (*34*) for the twenty-year periods centered on 2050 and 2070, under RCP 4.5 and RCP 8.5. We quantified exposure to climate change as the Euclidean distance between baseline and future climate values for each grid cell in a global Principal Component Analysis of eight bioclimatic variables (**table S3**). Then, based on global data, we calculated the 75^th^ percentile values of both stressors (exposure to land use impacts and climate change) for 2050 under RCP 8.5, thereby identifying VIPs with the highest exposure to these stressors individually and combined. (Different thresholds can be explored through the VIP-Explorer ShinyApp, **data S1**).

We found that under both RCPs ∼34-35% of VIPs occur within the quarter of the world’s terrestrial surface projected to experience the greatest exposure to land use impacts for 2050, with little net change projected for 2070. Most VIPs highly exposed to land use impacts are located within the Afrotropics, Neotropics and Indo-Malaya, and overwhelmingly within the Tropical and Subtropical Moist Broadleaf Forest, and Tropical and Subtropical Grassland, Savanna and Shrubland biomes (**Fig. 3, table S4-5**). In contrast, ∼13% of VIPs occur within the 25^th^ percentile of land projected to experience the lowest land use impacts by 2050. Hence, VIPs are over-represented throughout regions projected to have high exposure to land use impacts and under-represented in regions projected to have low land use impacts.

**Fig. 3.**
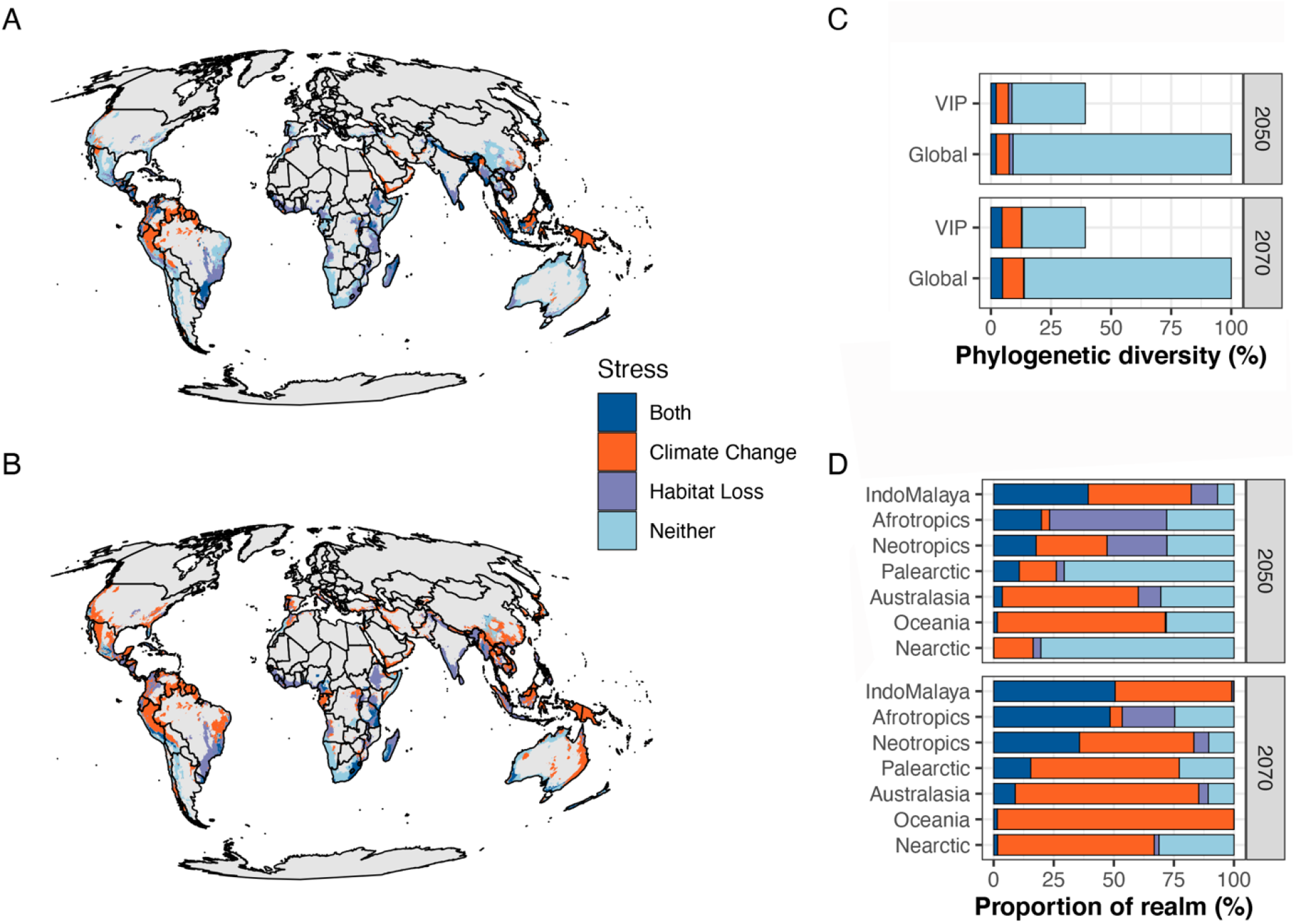
Exposure of hotspots of geographically restricted and evolutionarily distinct tetrapod lineages to land use impacts and climate change. **A**. The percentage of Phylogenetic Diversity either across the globe or unique to *Very Important Places* (VIPs) that is only found in grid cells with high exposure to (i) both land use impacts and climate change (dark blue), (ii) climate change only (orange), (iii) land use impacts only (lilac), or (iv) is not highly exposed to either stress (light blue). High exposure is defined as above the 75^th^ percentile of global values for 2050 under RCP 8.5. **B**. Proportion of VIPs within each biogeographic realm under high exposure to both, one or neither of the stresses. **C, D**. Location of VIPs under high exposure to both, one or neither of the stressors for 2050 (**C**) and 2070 (**D**).

We also found that by 2050, 25% (RCP 4.5) to 36% (RCP 8.5) of VIPs are projected to be located in areas with high exposure to climate change, increasing to 35% and 75% by 2070, respectively. High climate-exposed VIPs are concentrated in the Neotropical and Indo-Malayan realms, primarily within the Tropical and Subtropical Moist Broadleaf Forest biome (**table S6-7**). Again, for 2050 (RCP 8.5), VIPs are over-represented amongst the regions of the world projected to have high exposure to climate change.

While we did not directly assess the responses of tetrapods to either high exposure to land use impacts or climate change, these pressures will undoubtedly have serious consequences, particularly for species and lineages whose entire ranges are within highly-exposed VIPs. For 2050 (RCP 8.5), this includes 9% of global tetrapod evolutionary heritage, representing 3,891 species (**Fig. 3, fig. S2**). By 2070, this increases to 13% or 5,670 species, with almost all being highly exposed to climate change, with or without high exposure to land use impacts (**table S8**). In contrast, following a lower emissions trajectory (RCP 4.5), 8% of evolutionary heritage, representing 3,661 species, will be highly exposed by 2070, with almost all being highly exposed to climate change (**table S9**). Notably, when also considering areas other than VIPs, the proportion of global biodiversity highly exposed to these stressors does not substantially change, emphasizing the critical role of VIPs in representing global evolutionary heritage.

### Dual stressors will pose challenges to conservation efforts

VIPs where species are projected to experience high exposure to both stressors simultaneously pose a serious challenge to conservation. We call these VIPs the *wicked places*. For 2050 (RCP 8.5), ∼13% (n = 42,609) of VIPs are *wicked places*, predominantly occurring in Tropical and Subtropical Moist Broadleaf Forests and to a lesser extent, Tropical and Subtropical Grasslands, Savannas and Shrublands (66% and 12%, respectively). Almost three-quarters of these regions are confined to just 13 countries. For 2070, more than a quarter of VIPs are *wicked places* (29%, n = 95,648), with only 20% occurring outside of the Tropical and Subtropical biomes and <10% in the Nearctic and Palearctic realms (**Fig. 3**).

For countries harboring *wicked places*, we obtained the 2019 Readiness scores quantified by the Notre Dame Global Adaptation Initiative (ND-GAIN (*35*)) and the 2019 Human Development Index (HDI (*36*)), and calculated the global median of each. ND-GAIN assesses a country’s vulnerability to climate disruption as well as its readiness to adapt to climate change, measured as the ability of the country to leverage investments and convert these to adaptation actions. By 2050, over 90% of *wicked places* lie within countries with readiness scores below the global median. This includes India, Indonesia and Brazil which, combined, contain close to half of all *wicked places* (**Fig. 4A**). By 2070, the percent of *wicked places* in countries with readiness scores below the global median declines slightly (89% RCP 4.5; 83% RCP 8.5), although one quarter (RCP 4.5) to just over one third (RCP 8.5, **Fig. 4B**) lie in countries with scores within the lowest quartile (notably Ethiopia, Madagascar and Myanmar). Under RCP 8.5, more than two-thirds of *wicked places* (67.5%, 2050) lie within countries with HDI values below the global median, and although this decreases slightly to 62.3% for 2070 (**Fig. 4C**), more than one-third of *wicked places* (34.6%) are projected to be within countries with the lowest quartile HDI values (**Fig. 4D**).

**Fig. 4.**
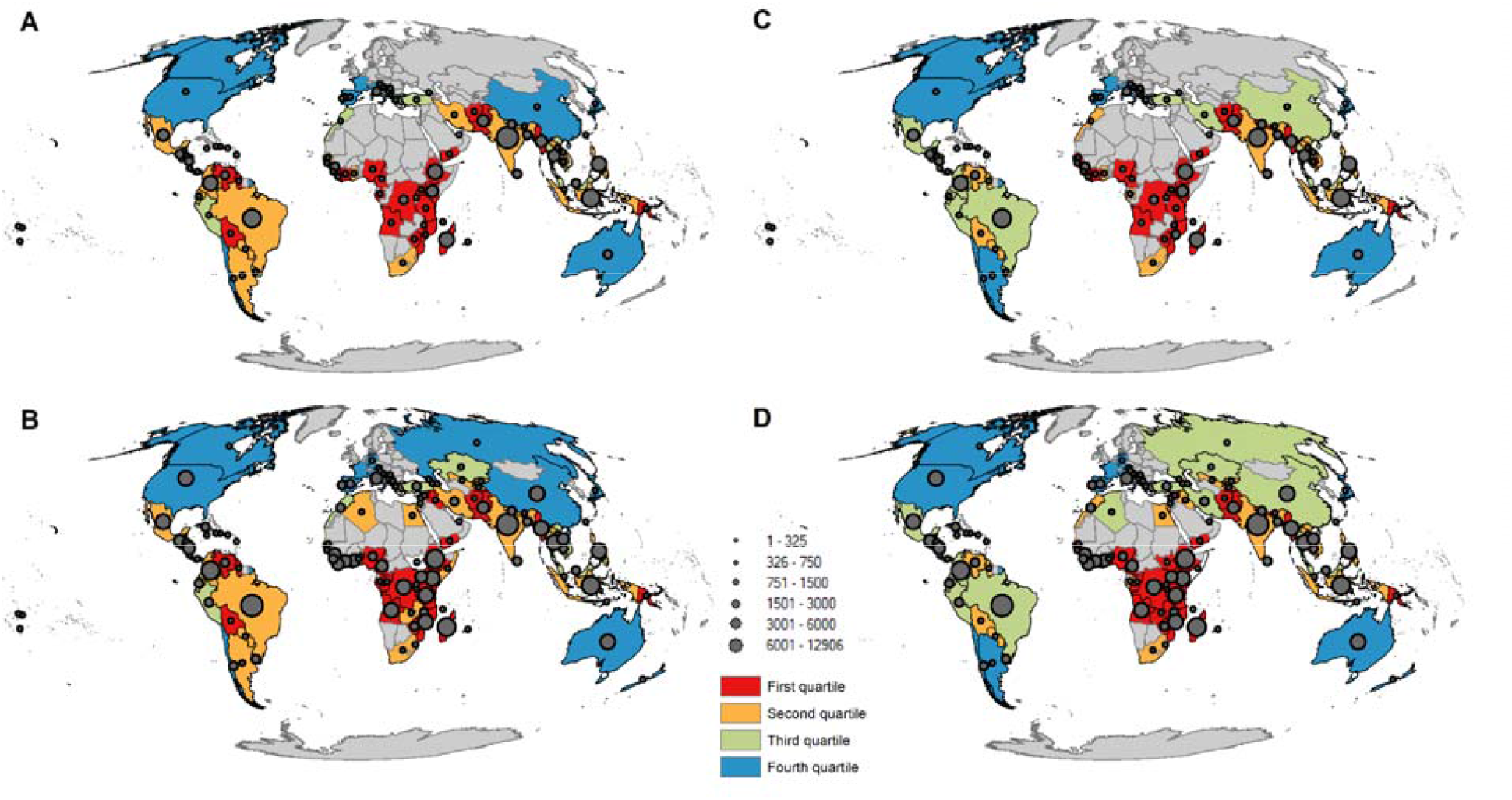
Relationship between area of w*icked places* in each country relative to country’s adaptive readiness score (A, B) and Human Development Index (C, D), for 2050 (A, C) and 2070 (B, D). *Wicked Places* represent hotspots of unique tetrapod evolutionary lineages projected to simultaneously face high exposure to land use impacts and climate change. The size of the circles represents the area encompassed by *wicked places* in each country. Colour of countries represents global quartiles for Readiness scores or Human Development Index. Countries without any *wicked places* are coloured grey.

## Discussion

The 2050 Vision for Biodiversity is one in which the world lives in harmony with nature (*4*). To achieve this vision, biodiversity loss must be halted by 2030, and ecosystem recovery over the following 20 years needs to be sufficient to result in net improvements to biodiversity outcomes by 2050 (*1*). Our study reveals important findings relevant to the 2050 Vision for Biodiversity. First, almost 40% of the world’s terrestrial tetrapod evolutionary heritage is restricted to one quarter of Earth’s land surface—regions we refer to as *Very Important Places* (VIPs). Second, under the business-as-usual scenario (RCP 8.5), these regions are over-represented amongst areas of the world predicted to experience high exposure to land use impacts or climate change by 2050. Third, of the VIPs projected to experience high magnitudes of both stressors simultaneously (the *wicked places*), the majority are located in countries with either an ND-GAIN Readiness score (indicating the country’s capacity to adapt to climate change) or human development index below the global median.

Identifying areas of highest exposure to both land use impacts and climate change provides a rapid means of locating regions in which biodiversity may experience the greatest pressure to undergo range shifts, while also being the most constrained from doing so (*37*). However, realized responses to these impacts will depend not just on their respective magnitudes, but also on species’ sensitivities (*38*) and the success of conservation actions and land use policies. By integrating predictions of exposure for two time periods (2050 and 2070), our analysis can be used to strategically identify actions required to minimize the long-term effects of land use impacts and climate change, and confer the greatest potential to secure species’ persistence and the evolutionary heritage they represent.

For VIPs in which exposure to land use impacts is predicted to be particularly high, protection and restoration will be crucial. Target 3 for Reducing Threats to Biodiversity in the Post-2020 Global Biodiversity Framework calls for ensuring that at least 30% of land area is conserved by 2030, including areas of importance to biodiversity (*1*). The spatial clustering of VIPs, and particularly that of *wicked places* in the Afrotropical and Indo-Malayan realms, provides support to calls for higher proportions of land to be protected in countries harboring rich or unique biota (*39*). However, area-based targets have the potential for perverse outcomes if they give rise to residual reservation with little benefit to biodiversity (*40*). Analyses using complementarity of Phylogenetic Diversity can aid with identifying investment priorities where conservation impact can be most effectively achieved.

While protection of VIPs should theoretically safe-guard their biodiversity, protected areas (PAs) do not necessarily experience lower human pressure than non-protected areas (*41*), particularly in Indo-Malaya, the Afrotropics and Neotropics where anthropogenic pressure on PAs has not abated in the last 15 years (*42*). For instance, non-forested PAs within tropical regions experience greater pressure from cropland conversion than in matched unprotected areas (*42*), and forested PAs continue to experience deforestation, albeit at lower rates than forests outside PAs (*43*). PAs require adequate resourcing if they are to be effective at halting biodiversity loss (*44, 45*), yet the effectiveness of PAs at limiting human pressure is positively associated with the human development index (*42*), for which six of the ten countries containing the greatest expanses of *wicked places* rank below the global median.

Large-scale restoration of ecosystems is critical to biodiversity conservation (*46*) and mitigation of climate change (*47*), with calls for 20% of degraded ecosystems to be restored by 2030 (*48*). Our analysis can help identify regions where restoration efforts may yield considerable benefits for the continuity of tetrapod evolutionary heritage. The Guinean forests of West Africa, Atlantic forests of South America, Irrawaddy region of southeast Asia and Ethiopian Highlands contain VIPs with high exposure to land use impacts. These regions have previously been identified as global priority areas for restoration, based on climate mitigation potential, cost minimization and benefits to biodiversity (*26*). However, these VIPs are also projected to be *wicked places* by 2070, highlighting the importance of using prospective-based—rather than retrospective— strategies (*49*) when undertaking restoration activities, to ensure resilience to anticipated climate change, such as the use of climate-ready germplasm (*50, 51*).

Indeed, under RCP 8.5, those areas most critical to conserving tetrapod evolutionary history are projected to become *wicked places* by 2070, facing the dual pressure of high exposure to land use impacts and climate change. Particularly problematic is that interaction between habitat loss and climate change can increase the impacts of land-cover change on birds and mammals by up to 43% and 24%, respectively (*52*). The adaptive capacity of biodiversity in VIPs highly exposed to climate change, either singularly or in combination with land use impacts, urgently needs to be assessed. While management of other threats may increase the survivability of populations in the face of climate change, habitat restoration and improvements to connectivity will facilitate dispersal of species across the landscape (*53*). For those species unable to undertake range shifts or to shift fast enough, identification and protection of refugia may be critical for their survival (*54*), while climate-sensitive species may require direct intervention through *ex situ* conservation, translocation or habitat engineering (*53, 54*).

We point out that VIPs with lower exposure to land use impacts and/or climate change will still require monitoring and improvement. Even in the absence of other types of threats, increases to the frequency and intensity of extreme events, such as heat waves, fires and droughts, superimposed on a warming climate may be sufficient to cause catastrophic ecosystem collapse, as has already been reported for diverse ecosystems (*55*), highlighting that we must expect the unexpected.

While we have mapped the overlap between tetrapod irreplaceability and exposure to two of the dominant threats to biodiversity for this century, there are some limitations to our study. First, expert-derived range maps may incorporate unsuitable habitat for the species in question (*56*) which, consequently, could under-estimate the pattern of irreplaceability. Conversely, irreplaceability could be over-estimated where range maps do not capture the entire realized distribution of the species. However, while there is some uncertainty in irreplaceability values per cell, our assessment has captured areas known for their high endemism (*24, 57*) of tetrapods (*58, 59*), suggesting that the use of expert range maps will not affect the broad sweep of our analysis.

Second, we considered only a very small proportion of the world’s biodiversity in our assessment. Tetrapods are commonly used in global analyses because of the availability and relative completeness of taxonomic, geographic and phylogenetic data, with the implicit assumption that observed patterns will be representative of broader biodiversity. However, the key drivers of patterns of endemism—geographic isolation and environmental gradients—apply to all taxa, with variation among groups being attributable to variation in dispersal capacity and environmental tolerance (*60*). Therefore, areas of conservation priority identified for one group should be broadly applicable to another, especially on a global scale (*61*).

We also point out that identifying the most irreplaceable regions and those with highest exposure to stress required that we set a threshold (here, at 75% of global scores). However, our framework can incorporate any threshold, which can then be used to direct local and regional management and conservation strategies, and alternative thresholds can be explored in our VIP-Explorer ShinyApp (**data S1**).

Rapidly reducing the deterioration of ecosystems, and conserving and restoring natural areas, has been cited as a highly cost-effective CO_2_ mitigation strategy that, if done wisely, can result in considerable co-benefits for wildlife (*62*). However, conservation is chronically underfunded, particularly in developing countries (*63*). Our findings highlight that the majority of the most irreplaceable regions for tetrapods are located in nations likely to lack resources to effectively manage these regions, and which have a higher proportion of their population dependent on nature to meet their daily needs (*64*). This presents major challenges, and avoiding future biodiversity losses will require unprecedented ambition and coordination as well as global pressure on governments to commit to agreements on climate finance, deforestation and emissions reductions made at Glasgow COP26.

## Author contributions

Conceptualization: LJB, DAN, PDW, JBB, MER

Methodology: LJB, DAN, PDW, JBB, MER

Investigation: LJB, DAN, PDW, JBB, MER

Visualization: LJB, DAN, PDW, JBB, MER

Writing - original draft: LJB, DAN, PDW, JBB, MER

Writing - review & editing: LJB, DAN, PDW, JBB, MER

## Competing interests

Authors declare that they have no competing interests.

## Data and materials availability

We used publicly available datasets for our analyses. Separate trees for birds (*65*), crocodiles and turtles (*66*), amphibians (*67*), mammals (*68*), and squamates (*69*) are available from sources. Species range data for mammals and amphibians are available from the IUCN RedList Portal (https://www.iucnredlist.org/resources/spatial-data-download) (*20*), bird data from BirdLife International and Handbook of the Birds of the World (http://datazone.birdlife.org/species/requestdis) (*21*), and reptiles from the Global Assessment of Reptile Distributions (http://www.gardinitiative.org/data.html) (*22*). Land-use projections were derived from the Land Use Harmonization project (https://luh.umd.edu/), while climate data were from CHELSA (https://chelsa-climate.org/). Most derived data are available through the VIP-Explorer ShinyApp (https://github.com/peterbat1/VIP_Explorer). Remaining data on the phylogenetic tree is available in SM, as is R code.

## Supplementary Materials

### Materials and Methods

#### Species range data

We included data on all tetrapods for which we could obtain expert-derived species range data. Our final list of species included 10,952 Aves (birds), 24 Crocodylia (crocodiles), 6,613 Lissamphibia (amphibians), 5,597 Mammalia (mammals), 1 Rhynchocephalia (tuatara), 9,690 Squamata (squamate reptiles), and 322 Testudines (turtles). Species range data were extracted from spatial polygon data acquired from three sources: (1) IUCN Red List of Threatened Species v6.2 (*20*) (marine mammals [last accessed 21 March 2019], terrestrial mammals [last accessed 2 December 2019], and amphibians [last accessed 2 December 2019]); (2) BirdLife International and Handbook of the Birds of the World (*21*) (last accessed 10 December 2019); and (3) the Global Assessment of Reptile Distributions v1.1 (GARD (*22*)). For birds, mammals and amphibians, we included range polygons with presence classified as “Extant” or “Probably Extant”, and origin classified as “Native”, “Reintroduced”, “Vagrant” (excluded for birds), “Origin Uncertain”, or “Assisted Colonisation”. Additionally, we included polygons with origin class “Introduced” if the species’ IUCN status was threatened. For birds, we excluded polygons with seasonality classified as “Passage”. We then manually excluded the ranges for species that were exclusively marine, retaining species with haul out sites.

Due to an error in the IUCN expert range for the frog *Litoria tyleri* (Pelodryadidae), we constructed a polygon to represent this species range. In doing so, we obtained occurrence records for *L. tyleri* from the Atlas of Living Australia (www.ala.org.au, last accessed 24 February 2021), and created a 10 km buffer around these points.

We converted all expert range polygons to global raster grids defined in the Mollweide coordinate system (ESRI:54009), at a resolution of 10 × 10 km. For each species, we identified all cells that overlapped range polygons (i.e., polygon covers cell center, or polygon vertex contained by cell). The subset of these cells that coincided with land (defined by the countries shapefile available at https://www.iucnredlist.org/resources/spatialtoolsanddata) was classified as the species’ range. After trimming out cells not defined as land, 33,199 tetrapod species and 1,340,520 grid cells were retained for further analysis.

#### Phylogenetic data

We compiled the phylogenetic tree of almost all known tetrapods (*n* = 33,199) by combining separate trees for birds (*65*), crocodiles and turtles (*66*), amphibians (*67*), mammals (*68*), and squamates (*69*). These trees were joined together by a tetrapod ‘backbone’ tree manually constructed following Crawford et al. (*70*) (**fig. S3**). The tuatara (*Sphenodon punctatus* [Sphenodontidae: Rhynchocephalia]) was included as a separate tip. Divergence dates between nodes of the tetrapod backbone tree were taken from timetree.org (*71*). The source tree for each major group of tetrapods was attached to the relevant branch of the backbone tree. The source trees included almost all known species and had branch lengths scaled relative to time since divergence (in millions of years). The source tree for crocodiles and turtles (*66*) spanned the clade Archosauromorpha, which includes birds as a single tip. We therefore extracted crocodile and turtle clades, and attached them separately to the backbone tree.

Species names in the source trees were matched to those used in the species range data. Where mismatches occurred, these were resolved on a case-by-case basis. For simple one-to-one synonyms and changes of genus, the name on our composite tetrapod tree was replaced with that associated with the species range data. For more complex cases, species names from the range data that were missing from the source tree were inserted into our composite tree within a clade defined by known close relatives (usually at the genus level). Exact placement within the clade was determined randomly (*72*). Details of decisions for all mismatched species names are in **data S2**.

Each of the source trees (except the tetrapod ‘backbone’ tree) were actually a set of a large number of versions of the tree, each version with differing topologies and divergence dates, representing uncertainty in phylogenetic relationships. For each source tree, we sampled 100 versions at random from the full set for our analyses, with the sample size of 100 being chosen based on computational constraints. For random insertions of missing species, this procedure was done independently for each version of the tree to model uncertainty as to the exact placement of the taxon.

#### Irreplaceability

We used complementarity of Phylogenetic Diversity (PD) as an indicator of the irreplaceability of a grid cell on a global scale. Complementarity in PD is defined as the sum of the branch lengths (in millions of years) of a phylogenetic tree that are unique to that grid cell (the focal cell) when compared to a subset of other cells (the comparison subset) (*23*). A branch is present in a grid cell if at least one of its descendent species occurs in the cell. Complementarity in PD contrasts with simple PD (*73*) in that it only sums the lengths of those branches that are present in the focal cell but absent from the comparison subset, while simple PD sums the lengths of all branches present in the focal cell, regardless of their presence or absence elsewhere. This measure captures information on the extent to which evolutionarily distinct lineages are spatially restricted to a focal cell, and is thus a form of phylogenetic endemism (*74*). Complementarity was calculated as an expected value when the focal cell is compared to 1000 other randomly selected cells (**fig. S4**). We then defined, and mapped, *Very Important Places* (VIPs) as cells with a complementarity value ≥ the 75^th^ percentile, based on all terrestrial cells.

Rather than estimating a mean from repeated Monte Carlo sampling of 1000 cells from the dataset, expected complementarity values were determined from an exact analytical equation (*19*), which calculates the mean complementarity across all possible combinations of 1000 cells from all terrestrial cells (*n* = 1,340,520). Given a phylogenetic tree with *T* branches, each with a length of *l*_*k*_, and a total of *n* cells, from which a random subset of size *m* is drawn, the expected complementarity of a focal cell is calculated from Equation 1.

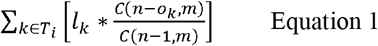

The fraction gives the probability that a particular branch (*k*) of the phylogenetic tree that is present in the focal cell, is absent from a random selection of *m* other cells. The numerator is the number of possible combinations of *m* cells where branch *k* is absent, and the denominator is the total number of possible combinations of *m* cells that does not include the focal cell. For any given subset size of *m* randomly chosen cells, the probability is determined by the total number of grid cells (*n*) and the number of grid cells occupied by the branch (*o*_*k*_). Multiplying the probability of the branch (being absent) by its length and then summing across the set of branches present in the focal cell (*T*_*i*_) gives the expected PD uniquely found in that cell. Calculations were performed for each of 100 versions of the tetrapod phylogenetic tree and then averaged.

Diagnostic analysis showed that a comparison subset size (*m*) of 1000 produced complementarity values that were not strongly correlated with simple PD (Pearson’s *r* = 0.28), ensuring that the measure captured information on endemism more than alpha diversity (**fig. S5**). While larger subset sizes had lower correlations with alpha diversity, complementarity values of all cells converge towards zero as subset sizes increase (**fig. S4**). This is because, at this spatial resolution, very few species or branches occur in only a single cell, and thus more and more grid cells drop to zero complementarity as subset sizes approach the total number of available grid cells. The subset size of 1000 chosen for this study strikes a good balance between avoiding being overly correlated with alpha diversity while also not employing an overly strict definition of ‘endemism’ with minimal variance among cells.

Complementarity in PD was also compared to complementarity in species richness, as well as species richness (i.e., count of species) and simple PD for each grid cell. Complementarity in species richness is defined as the count of species found in a focal cell but absent from the comparison subset (of size 1000). The measure was also calculated from an exact analytical equation (Equation 2). For a set of species found in a focal cell (*S*_*i*_), the probability of each of those species being absent from a randomly selected subset of *m* other cells depends on the total number of grid cells (*n*) and the number of grid cells occupied by each species (*o*_*j*_). Correlations (Pearson) between each measure for all tetrapods and for all major groups (i.e., amphibians, mammals, squamates, turtles, crocodilians and birds) were calculated for all pairwise comparisons (**data S3**).

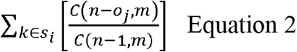

#### Land-Use Harmonization (LUH2) data

Changes in land use were described by the Land-Use Harmonization (LUH2) datasets (version 2f) developed as part of the Land-Use Model Intercomparison Project (LUMIP) (*32*). These datasets reflect a set of land-use scenarios spanning historical reconstructions of land-use through to future projections, and include data for the years 850–2100. Future states (2015–2100) use the sixth phase of the Coupled Model Intercomparison Project (CMIP6) future scenarios at a spatial resolution of 0.25º x 0.25º. Data comprise 12 land-use categories, including primary and secondary forest and non-forest sub-types, managed pasture and rangeland, and five crop functional types. We downloaded future data, in NetCDF format, corresponding to SSP2 RCP4.5 (from MESSAGE-GLOBIOM) and SSP5 RCP8.5 (from ReMIND-MAgPIE) (http://luh.umd.edu/data.shtml). For these land-use change scenarios, we extracted the proportion of each grid cell covered by each of the 12 land-use types for the baseline period (2015), as well as for 2050 and 2070. We then multiplied these proportions by coefficients describing the relationship between land-use class, and whether a cell is forested or unforested, to the overall habitat condition of land at a cell (*33*) (**table S2**). The coefficients reflect the number of vascular plant species expected to occur within a given land-use, proportional to the number expected to occur if the environment was pristine. We then subtracted the value of each grid cell from 1 to represent the loss in habitat condition arising from land use impacts. The resulting layers (2015, and RCP 4.5 and RCP 8.5 for both 2050 and 2070 **fig. S6**) were then reprojected to a 10 × 10 km resolution grid in the Mollweide projection (ESRI:54009) using GDAL (www.gdal.org) with the R package gdalUtils (*75*).

#### Climate data

To quantify projected exposure to climate change, we used global climate data provided by the CHELSA v1.2 data set (*34*). These data provide a climate baseline from 1979–2013, at a spatial resolution of 30 arc seconds (approximately 1 km at the equator), and were created by interpolation methods designed to accurately represent the effects of topography, humidity, wind strength and direction on temperature and precipitation. A detailed description of the generation of these data is given elsewhere (*34, 76*). CHELSA contains a set of 19 bioclimatic variables (BIOCLIM) derived from monthly mean, maximum and minimum temperature and mean precipitation values, which are frequently used in ecological applications. Of these, we selected a subset of eight that represent annual average and seasonal contrasts for temperature and precipitation (*77*): annual mean temperature, temperature seasonality, mean temperature of the warmest and coldest quarters, annual precipitation, precipitation seasonality, and precipitation in the wettest and driest quarters (**table S3**). These data were downloaded from the CHELSA v1.2 repository (www.chelsa.org, last accessed 11 November 2020) using R scripts (see **code S1**).

The CHELSA dataset also provides downscaled output from a broad selection of future climate models from the Climate Model Inter-comparison Project v5 (CMIP5) (*78*), which formed the basis of the Inter-governmental Panel on Climate Change (IPCC) Fifth Assessment Report (*79*). The downscaling method, described by Karger et al. (*34*) applies the same technique used for interpolation of observed climate. Using model performance rankings (*80, 81*), we selected the top 15 GCMs (**table S10**) to represent the range of likely future climate conditions. We used the moderate and high emissions concentration pathways, RCP 4.5 and RCP 8.5 (*82*), respectively, to represent the most likely span of possible climate outcomes. Future climate was represented by mean conditions centered on 2050 (2041–2060) and 2070 (2061–2080) as supplied in the CHELSA dataset. Similar to baseline climate data, data describing future values for the selected eight bioclim variables were downloaded using R scripts (see **code S1**).

We projected the fine-resolution climate grids onto a Mollweide equal area projection (ESRI:54009) with 10 × 10 km grid cells, with the value assigned being the average from the finer resolution grid. All data were then assembled into a single data table consisting of values of each bioclimatic variable in the baseline climate, and 60 rows (for the GCM*RCP*future epoch combinations, i.e.,15 GCMs, two RCPs, two future epochs), for each of the eight bioclimatic variables. A Principal Component Analysis (PCA) was performed on the pooled data, which ensured that the scaling of data was uniform across grid cells and the climate exposure values computed in PCA space would be comparable.

We defined climate exposure as the Euclidean distance in PCA space between the baseline conditions for a grid cell and future conditions. Distances in transformed spaces have previously been used to characterize climate exposure (*83, 84*). Our method, similar to a method applied to model performance ranking (*81*), ensures that correlations between climate variables are accounted for and that the same transformation is applied to all data.

Finally, for each combination of RCP and future epoch, we calculated the mean distance across the 15 GCMs, resulting in four exposure values for each cell (i.e., two RCP*two epochs) (**fig. S7**).

#### Exposure to land use and climate change

For both stressors, land use impacts and climate change, we quantified the 75^th^ percentile value across all grid cells under RCP 8.5 for 2050, classifying areas above this threshold as ‘high exposure’. Using this threshold for both scenarios and future time periods enabled all results to be compared to the same standard. We then assessed the congruence between VIPs and the two stressors, defining as *wicked places* those VIPs projected to be simultaneously exposed to high values of both stressors. We summarized the number of VIP cells across biogeographic realms and biomes, based on the Terrestrial Ecoregions of the World (*85*). We downloaded the University of Notre Dame Global Adaptation Index (*35*) (ND-GAIN), and calculated the global median Readiness value for 2019. Readiness represents the capacity of a country to leverage both private and public investments to undertake climate change adaptation actions. We applied the same approach to the Human Development Index (*36*) (HDI), and calculated the percentage of *wicked places* that lie within countries with ND-GAIN or HDI below the global median.

Those species and lineages whose entire ranges were only found within regions of high stress (≥ 75^th^ percentile) for one or both stressors were considered to be ‘highly exposed’ and counted towards the proportion of biodiversity most at risk (**Fig. 3**). The proportion of species richness that is highly exposed is simply derived from the count of species whose entire ranges fall within regions of high stress. The proportion of phylogenetic diversity highly exposed was calculated as the sum of the lengths of branches in the phylogenetic tree whose ranges were entirely within regions of high stress. The range of a branch is the set of cells occupied by one or more of the species descended from that branch. By only counting species and branches whose ranges were wholly exposed to high stress, it was possible to additively partition biodiversity into those components exposed to: both land use impacts and climate change; climate change only; land use impacts only; or not highly exposed to either stress. Calculations were done for all species and lineages, as well as for that subset that are uniquely found in VIPs.

**Fig. S1.**
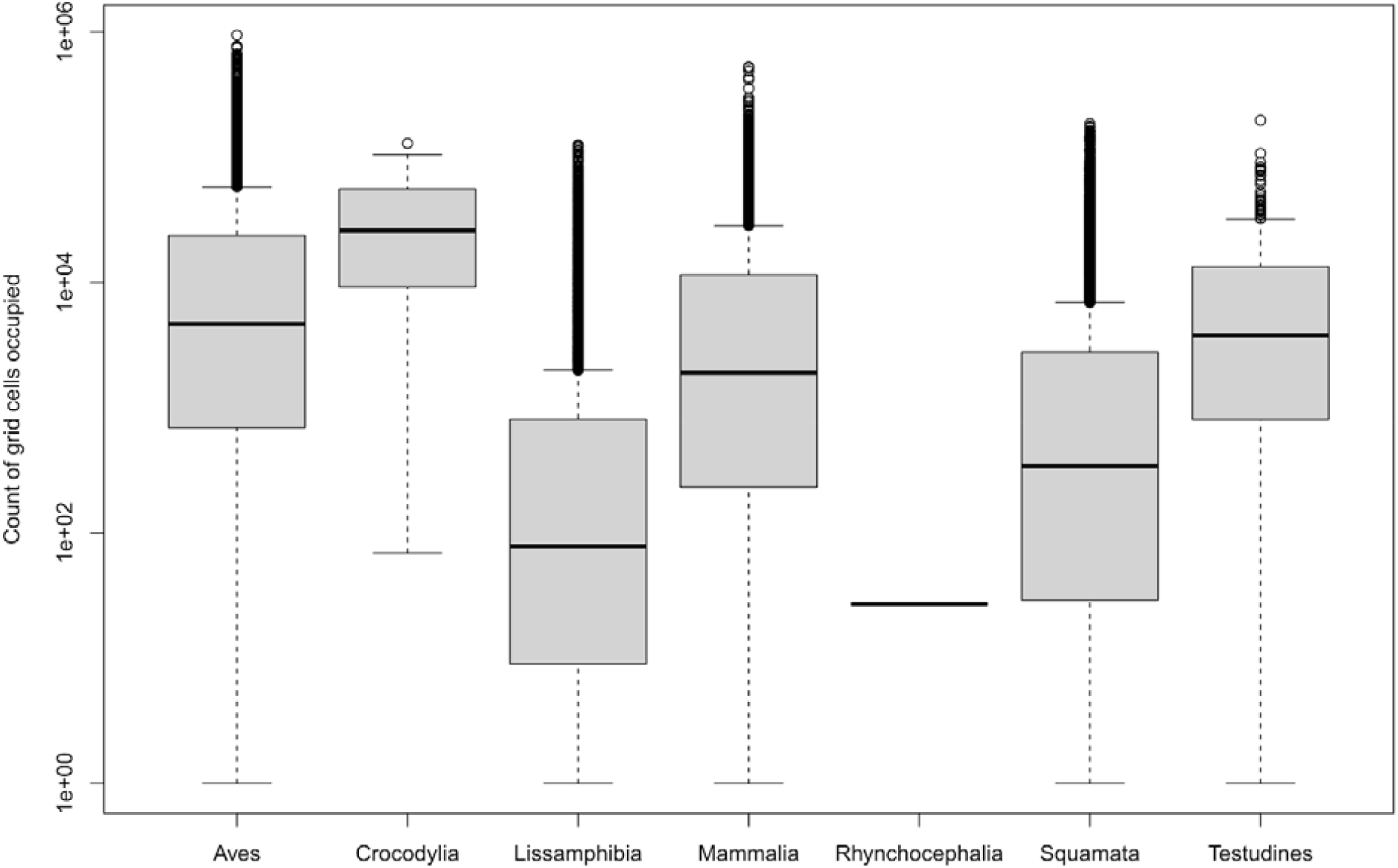
Distribution of range sizes (as count of occupied 10 × 10 km grid cells) for each major group of tetrapods. Box and whisker plot shows median and interquartile range (as grey box). Outliers (more than 1.5x upper quartile) are shown as individual points.

**Fig. S2.**
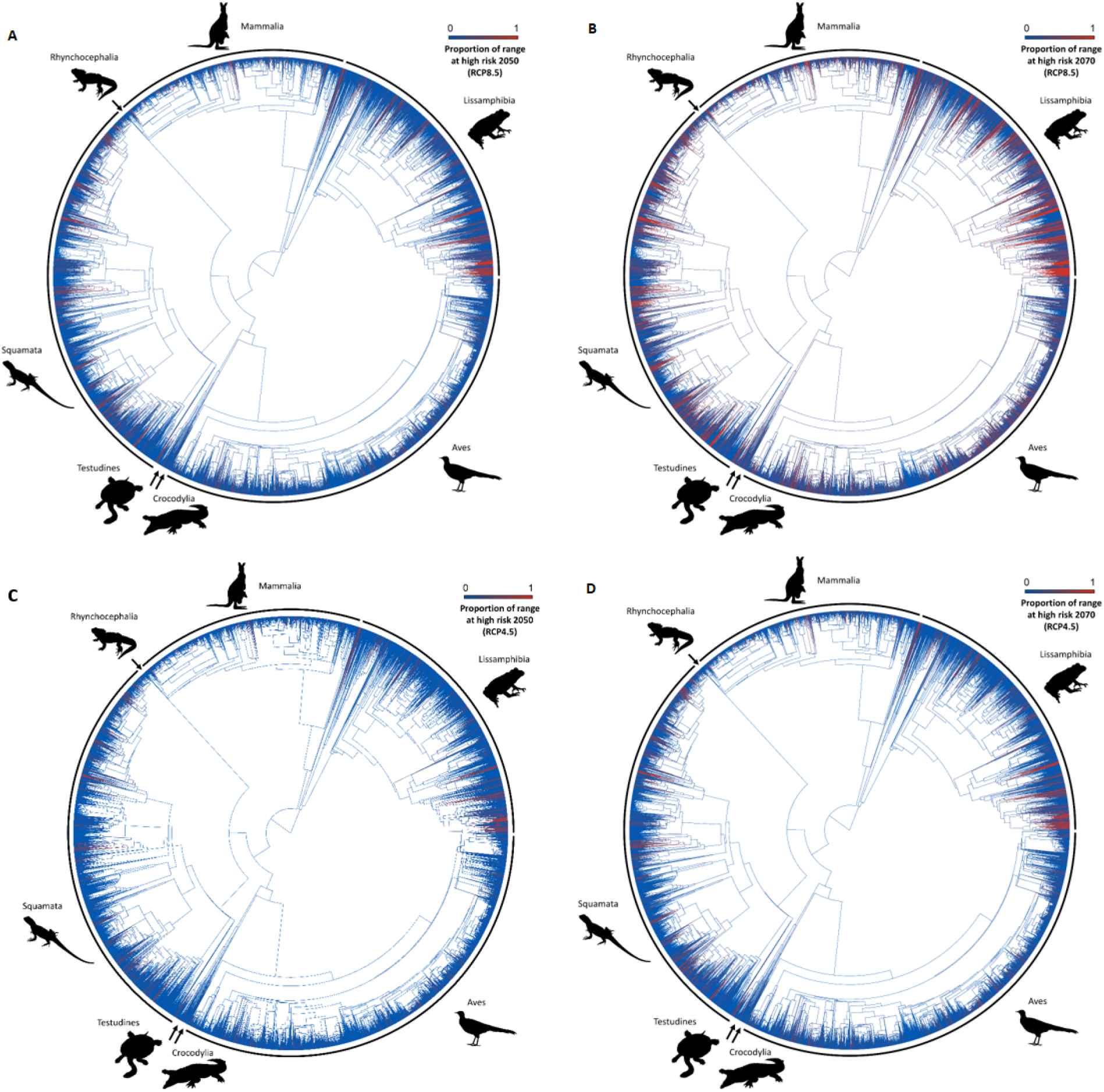
Phylogenetic tree of tetrapod species. Phylogenetic tree of 33,199 tetrapod species (mammals [Mammalia], birds [Aves], squamate reptiles [Squamata], crocodiles [Crocodylia], turtles [Testudines], tuatara [Rhynchocephalia] and amphibians [Lissamphibia]). Red branches represent those that are restricted to regions defined as *Very Important Place*s (VIPs) (i.e., 25% of the terrestrial surface with the highest concentration of geographically restricted and evolutionarily distinct tetrapod lineages) and which are at high risk of both exposure to climate change and land use impact by 2050 **(A, C)** or 2070 **(B, D)**, under Representative Concentration Pathway 8.5 **(A, B)** or 4.5 **(C, D)**.

**Fig. S3.**
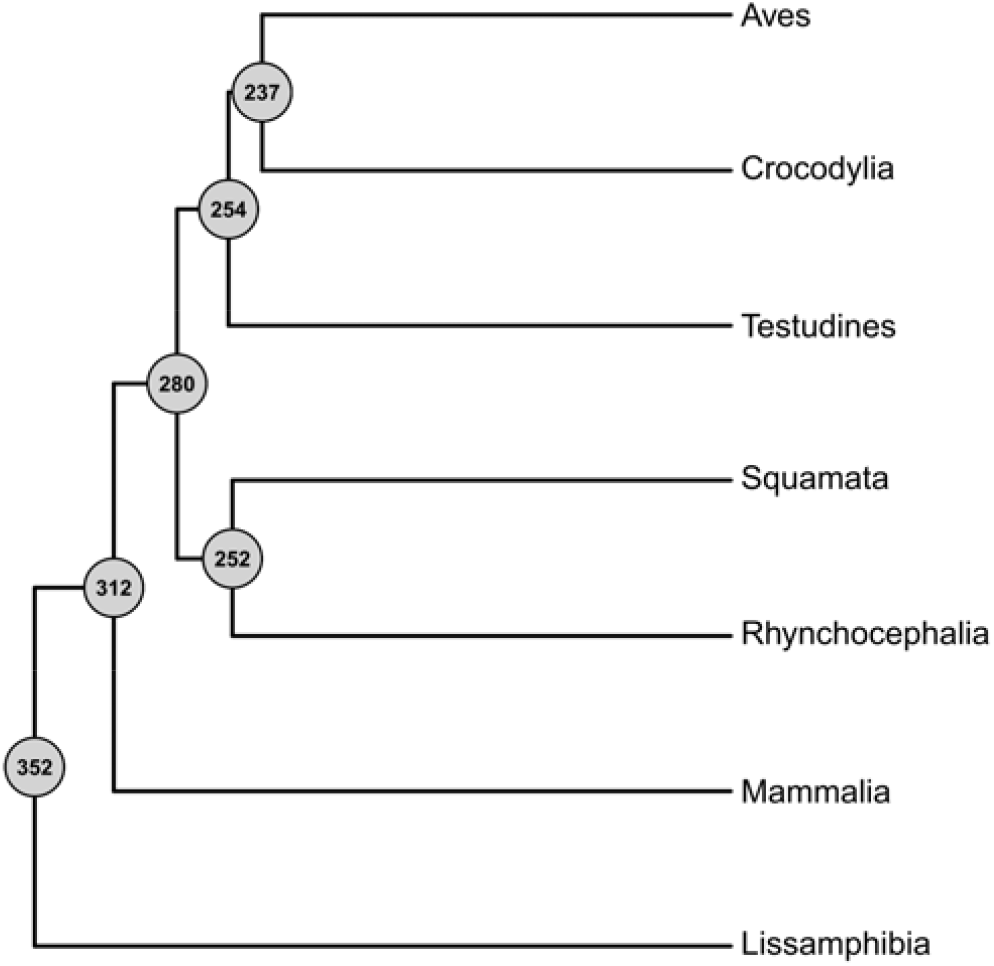
Phylogenetic tree depicting divergences between the major clades of tetrapods with divergence dates (Ma) in circles at each node. The tree was used as a backbone connecting separate phylogenetic trees of amphibians (Lissamphibia), mammals (Mammalia), squamate (Squamata) reptiles, turtles (Testudines), crocodiles (Crocodylia) and birds (Aves). The topology of the tree follows (*70*). Divergence dates are mean estimates sourced from timetree.org (*71*).

**Fig. S4.**
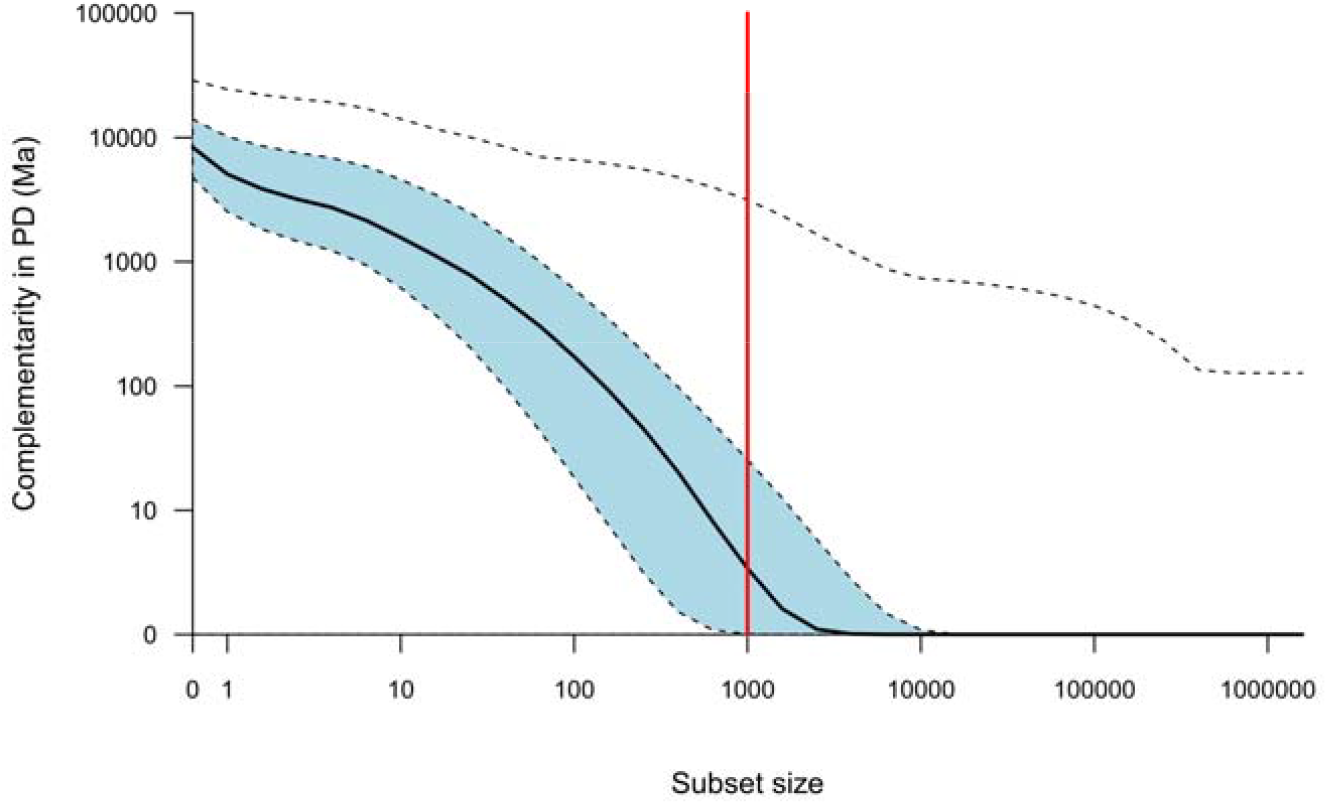
Complementarity in the Phylogenetic Diversity (PD) of terrestrial tetrapods for all 10 × 10 km grid cells. Complementarity is calculated as the PD uniquely represented in a cell when compared to different sizes of subsets of randomly selected cells. The solid black line is the median for all cells. The blue shaded area depicts the interquartile range. The uppermost dashed line is the maximum value observed. The red vertical line marks the subset size chosen for this study. This diagnostic analysis used a single version of the tetrapod tree with branch lengths scaled to millions of years (Ma) between nodes.

**Fig. S5.**
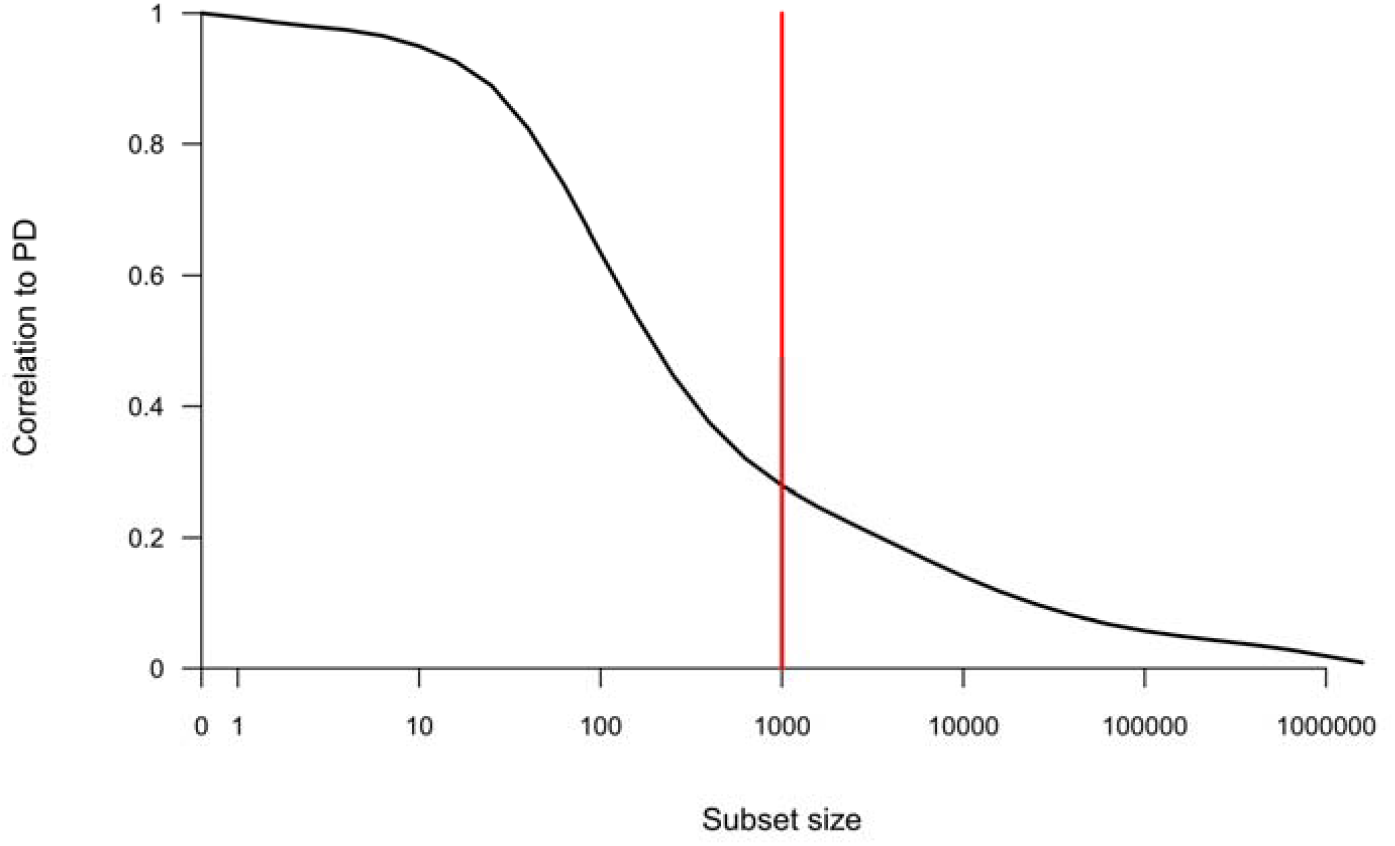
Correlation (Pearson) between Phylogenetic Diversity (PD) and complementarity in Phylogenetic Diversity of terrestrial tetrapods for all 10 × 10 km grid cells. Complementarity is calculated as the PD uniquely represented in a cell when compared to different sizes of subsets of randomly selected cells. The red vertical line marks the subset size (1000) chosen for this study. This diagnostic analysis used a single version of the tetrapod tree.

**Fig. S6.**
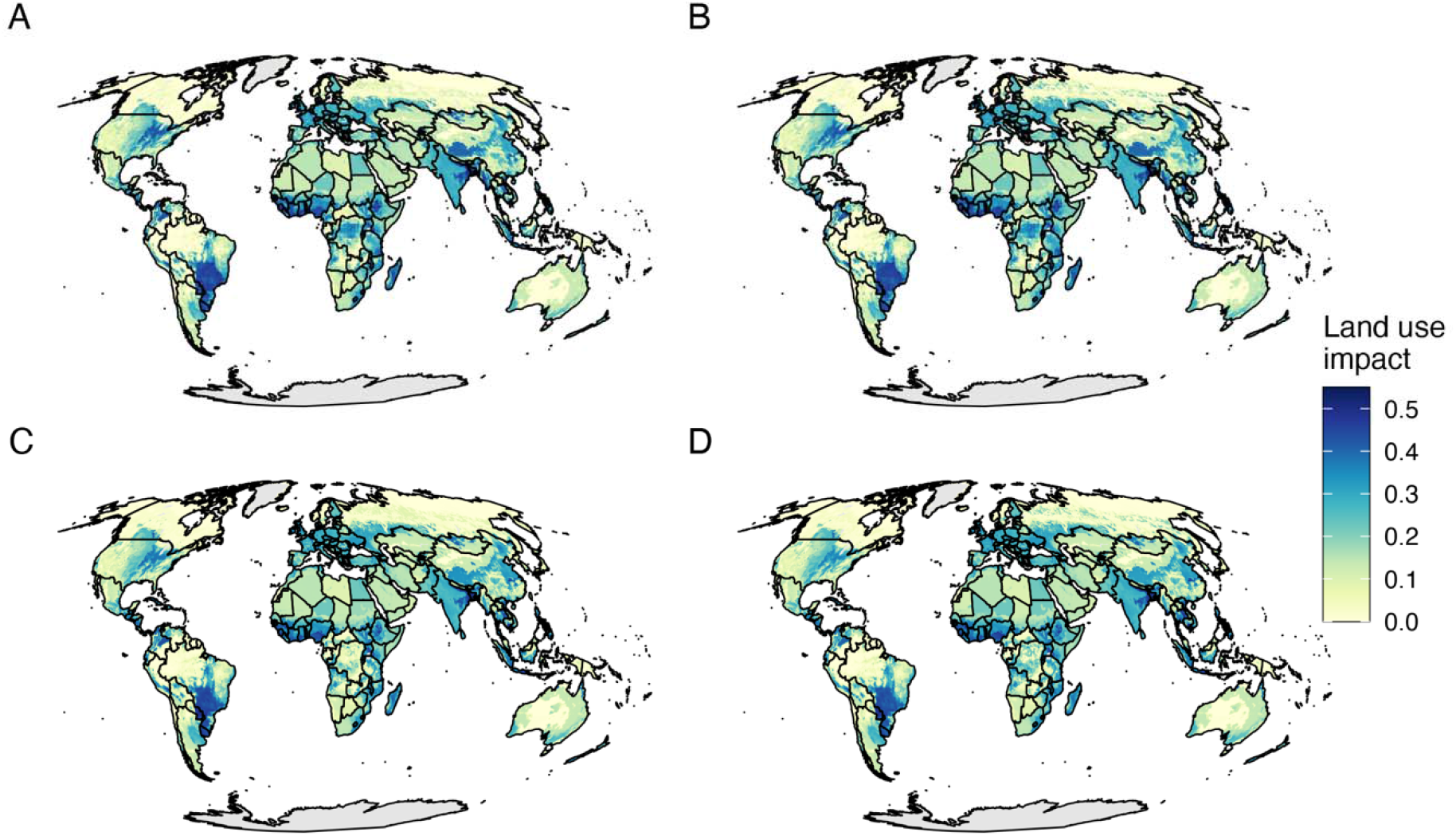
Projected land use impact. Land use impact is reflective of the number of species that may be lost from an area, depending upon projected changes to 12 types of land uses. Maps are for 2050 **(A, B)** and 2070 **(B, D)**, under Representative Concentration Pathway 8.5 **(A, B)** or 4.5 **(C, D)**.

**Fig. S7.**
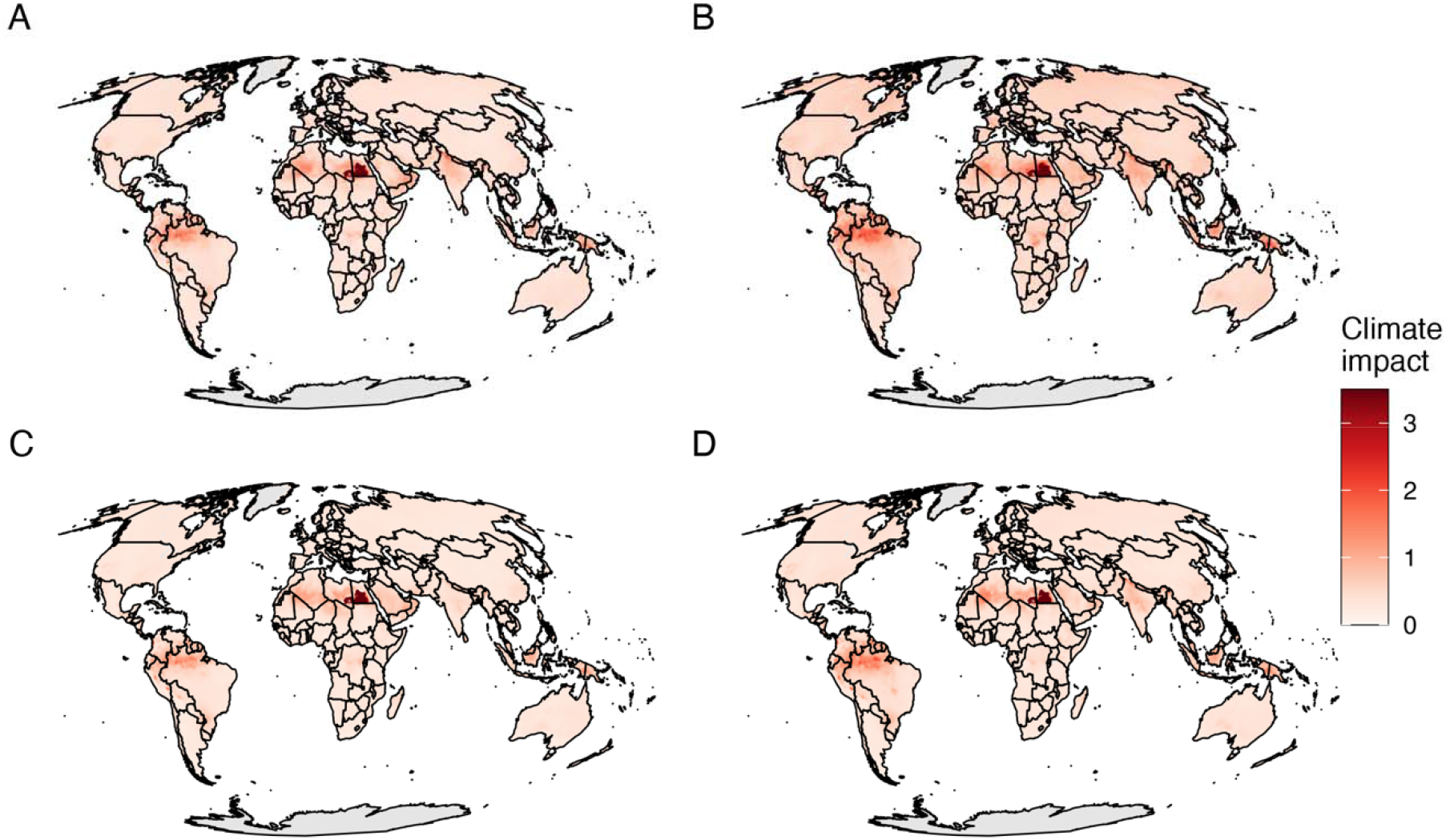
Projected exposure to climate change. This was calculated as Euclidean distance in principal component space based on eight bioclimatic variables (**table s4**), and averaged across 15 Global Climate Model scenarios for twenty-year periods centred on 2050 **(A, C)** and 2070 **(B, D)**, under Representative Concentration Pathway 8.5 **(A, B)** or 4.5 **(C, D)**.

**Table S1.**
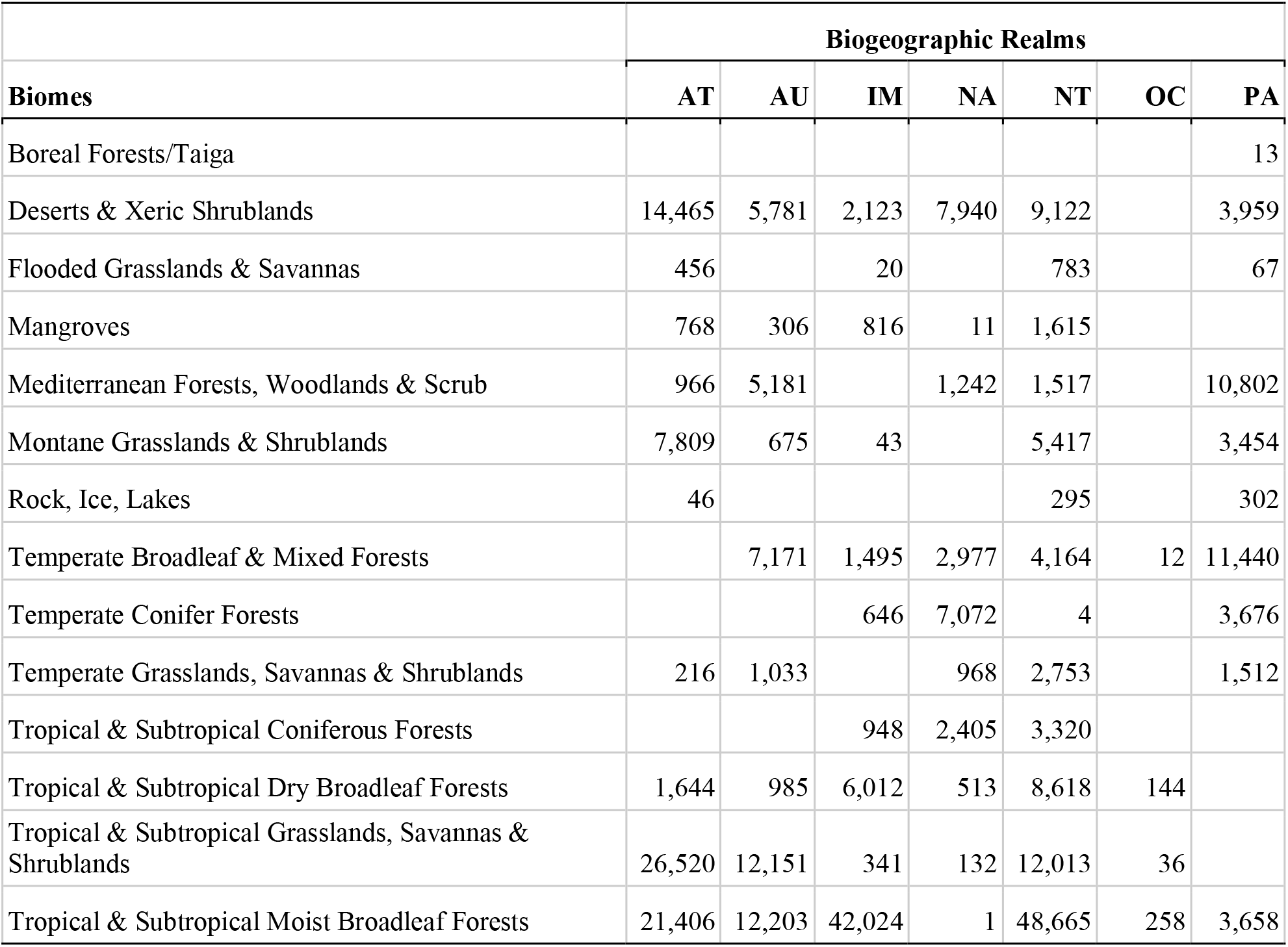
Number of VIP cells within each realm and biome. VIP refers to *Very Important Places*, which we define as the quarter of the Earth’s terrestrial area that has the highest irreplaceability scores for tetrapods, based on 10 × 10 km grid cells, hence VIPs = 335,130 grid cells. Realms and bioregions are from Terrestrial Ecoregions of the World Version 2.0 (*85*). AT = Afrotropics, AU = Australasia, IM = Indo-Malaya, NA = Nearctic, NT = Neotropics, OC = Oceania, PA = Palearctic.

| Biomes | Biogeographic Realms |  |  |  |  |  |  |
| --- | --- | --- | --- | --- | --- | --- | --- |
|  | AT | AU | IM | NA | NT | OC | PA |
| Boreal Forests/Taiga |  |  |  |  |  |  | 13 |
| Deserts & Xeric Shrublands | 14,465 | 5,781 | 2,123 | 7,940 | 9,122 |  | 3,959 |
| Flooded Grasslands & Savannas | 456 |  | 20 |  | 783 |  | 67 |
| Mangroves | 768 | 306 | 816 | 11 | 1,615 |  |  |
| Mediterranean Forests, Woodlands & Scrub | 966 | 5,181 |  | 1,242 | 1,517 |  | 10,802 |
| Montane Grasslands & Shrublands | 7,809 | 675 | 43 |  | 5,417 |  | 3,454 |
| Rock, Ice, Lakes | 46 |  |  |  | 295 |  | 302 |
| Temperate Broadleaf & Mixed Forests |  | 7,171 | 1,495 | 2,977 | 4,164 | 12 | 11,440 |
| Temperate Conifer Forests |  |  | 646 | 7,072 | 4 |  | 3,676 |
| Temperate Grasslands, Savannas & Shrublands | 216 | 1,033 |  | 968 | 2,753 |  | 1,512 |
| Tropical & Subtropical Coniferous Forests |  |  | 948 | 2,405 | 3,320 |  |  |
| Tropical & Subtropical Dry Broadleaf Forests | 1,644 | 985 | 6,012 | 513 | 8,618 | 144 |  |
| Tropical & Subtropical Grasslands, Savannas & Shrublands | 26,520 | 12,151 | 341 | 132 | 12,013 | 36 |  |
| Tropical & Subtropical Moist Broadleaf Forests | 21,406 | 12,203 | 42,024 | 1 | 48,665 | 258 | 3,658 |

**Table S2.** Coefficients used to convert land-use categories into habitat condition scores. From Di Marco et al. (*36*)

| Land-use | Forested biomes | Non-forested biomes |
| --- | --- | --- |
| Primary vegetation (forest and non-forest) | 1 | 1 |
| Secondary vegetation (forest and non-forest) |  |  |
| Mature ($\geq 50$ yrs) | 0.990 | 0.846 |
| Intermediate (30–50 yrs) | 0.855 | 0.883 |
| Young ($< 30$ yrs) | 0.757 | 0.778 |
| Perennial croplands (C3 and C4) | 0.635 | 0.687 |
| Nitrogen-fixing croplands | 0.585 | 0.749 |
| Annual croplands (C3 and C4) | 0.460 | 0.684 |
| Rangelands | 0.550 | 0.863 |
| Managed pasture | 0.533 | 0.587 |
| Urban | 0.645 | 0.717 |

**Table S3.** Eight bioclimatic variables used to quantify climate stress.

| **Bioclim variable** | **Definition (units)** | **Description** |
| --- | --- | --- |
| bio01 | Annual mean temperature (Celsius) | The mean temperature of a 12 monthly period. Calculated as the mean of the summed average temperature of the *i*th day of a year. |
| bio04 | Temperature seasonality (Celsius, standard deviation of mean monthly temperatures) | The amount of temperature variation observed over a given year, calculated using the coefficient of variation. Values are converted to degrees Kelvin (i.e. 273.15) to avoid dividing by zero. |
| bio10 | Mean temperature of the warmest quarter (Celsius) | The mean temperature during the quarter with the warmest temperatures. A quarter is defined as three consecutive months (1,2,3,...,10,11,12). |
| bio11 | Mean temperature of the coldest quarter (Celsius) | The mean temperature during the quarter with the coldest temperatures. A quarter is defined as three consecutive months (1,2,3,...,10,11,12). |
| bio12 | Annual rainfall (mm) | The sum of all precipitation values. |
| bio15 | Precipitation seasonality (dimensionless, coefficient of variation) | The amount of total monthly precipitation varying over a given year, calculated using the coefficient of variation. |
| bio16 | Precipitation in wettest quarter (mm) | The maximum total precipitation during a quarter. A quarter is defined as three consecutive months (1,2,3,...,10,11,12). |
| bio17 | Precipitation in the driest quarter (mm) | The minimum total precipitation during a quarter. A quarter is defined as three consecutive months (1,2,3,...,10,11,12). |

**Table S4.**
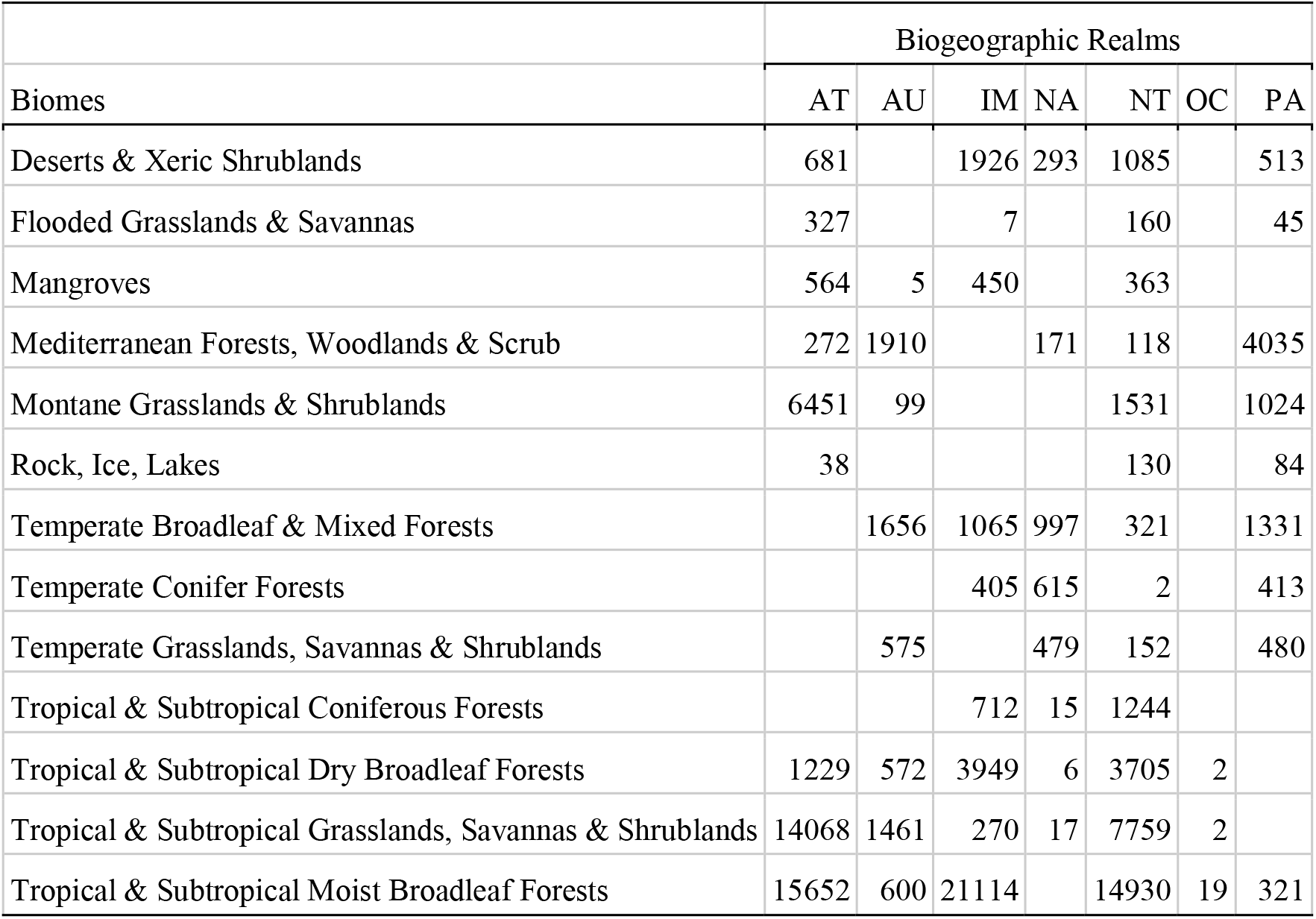
Number of VIP cells highly exposed to land use impacts for 2070, within each realm and biome, under scenario RCP 8.5. ‘Highly exposed’ is defined as a quarter of the Earth’s terrestrial surface with a loss of habitat condition value exceeding the 75^th^ percentile threshold for 2050, under Representative Concentration Pathway 8.5. *Very Important Places* (VIPs) span a quarter of the Earth’s terrestrial area that has the highest irreplaceability scores for tetrapods, based on 10 × 10 km grid cells, hence VIPs = 335,130 grid cells. Realms and bioregions are from Terrestrial Ecoregions of the World Version 2.0 (*85*). AT = Afrotropics, AU = Australasia, IM = Indo-Malaya, NA = Nearctic, NT = Neotropics, OC = Oceania, PA = Palearctic.

**Table S5.** Number of VIP cells highly exposed to land use impacts for 2070, within each realm and biome, under scenario RCP 4.5. ‘Highly exposed’ is defined as a quarter of the Earth’s terrestrial surface with a loss of habitat condition value exceeding the 75^th^ percentile threshold for 2070, under Representative Concentration Pathway 8.5. *Very Important Places* (VIPs) span a quarter of the Earth’s terrestrial area that has the highest irreplaceability scores for tetrapods, based on 10 × 10 km grid cells, hence VIPs = 335,130 grid cells. Realms and bioregions are from Terrestrial Ecoregions of the World Version 2.0 (*85*). AT = Afrotropics, AU = Australasia, IM = Indo-Malaya, NA = Nearctic, NT = Neotropics, OC = Oceania, PA = Palearctic.

| Biomes | Biogeographic Realms |  |  |  |  |  |  |
| --- | --- | --- | --- | --- | --- | --- | --- |
|  | AT | AU | IM | NA | NT | OC | PA |
| Deserts & Xeric Shrublands | 763 | 3 | 1854 | 300 | 1223 |  | 487 |
| Flooded Grasslands & Savannas | 261 |  | 2 |  | 231 |  | 45 |
| Mangroves | 535 | 7 | 484 |  | 392 |  |  |
| Mediterranean Forests, Woodlands & Scrub | 185 | 1916 |  | 114 | 118 |  | 5497 |
| Montane Grasslands & Shrublands | 6135 | 95 |  |  | 1544 |  | 1174 |
| Rock, Ice, Lakes | 37 |  |  |  | 132 |  | 97 |
| Temperate Broadleaf & Mixed Forests |  | 1532 | 1040 | 462 | 340 |  | 2044 |
| Temperate Conifer Forests |  |  | 354 | 253 | 2 |  | 515 |
| Temperate Grasslands, Savannas & Shrublands |  | 605 |  | 486 | 205 |  | 592 |
| Tropical & Subtropical Coniferous Forests |  |  | 632 | 12 | 1326 |  |  |
| Tropical & Subtropical Dry Broadleaf Forests | 1150 | 763 | 4521 | 5 | 3806 | 2 |  |
| Tropical & Subtropical Grasslands, Savannas & Shrublands | 13188 | 1500 | 243 | 18 | 7469 |  |  |
| Tropical & Subtropical Moist Broadleaf Forests | 13414 | 920 | 21279 |  | 13235 | 14 | 321 |

**Table S6.** Number of VIP cells highly exposed to climate change for 2070, within each realm and biome, under scenario RCP 8.5. ‘Highly exposed’ is defined as a quarter of the Earth’s terrestrial surface with a loss of habitat condition value exceeding the 75^th^ percentile threshold for 2050, under Representative Concentration Pathway 8.5. *Very Important Places* (VIPs) span a quarter of the Earth’s terrestrial area that has the highest irreplaceability scores for tetrapods, based on 10 × 10 km grid cells, hence VIPs = 335,130 grid cells. Realms and bioregions are from Terrestrial Ecoregions of the World Version 2.0 (*85*). AT = Afrotropics, AU = Australasia, IM = Indo-Malaya, NA = Nearctic, NT = Neotropics, OC = Oceania, PA = Palearctic.

| Biomes | Biogeographic Realms |  |  |  |  |  |  |
| --- | --- | --- | --- | --- | --- | --- | --- |
|  | AT | AU | IM | NA | NT | OC | PA |
| Boreal Forests/Taiga |  |  |  |  |  |  | 13 |
| Deserts & Xeric Shrublands | 4246 | 2182 | 2028 | 6532 | 6083 |  | 3226 |
| Flooded Grasslands & Savannas | 337 |  | 17 |  | 461 |  | 45 |
| Mangroves | 655 | 306 | 777 | 8 | 1200 |  |  |
| Mediterranean Forests, Woodlands & Scrub | 3 | 398 |  | 762 | 473 |  | 8476 |
| Montane Grasslands & Shrublands | 3747 | 329 | 43 |  | 962 |  | 2806 |
| Rock, Ice, Lakes | 39 |  |  |  | 134 |  | 301 |
| Temperate Broadleaf & Mixed Forests |  | 3416 | 1492 | 2977 | 2852 |  | 9691 |
| Temperate Conifer Forests |  |  | 641 | 5942 |  |  | 2391 |
| Temperate Grasslands, Savannas & Shrublands | 216 | 241 |  | 665 | 160 |  | 1342 |
| Tropical & Subtropical Coniferous Forests |  |  | 948 | 2066 | 2220 |  |  |
| Tropical & Subtropical Dry Broadleaf Forests | 1175 | 733 | 5728 | 403 | 5739 | 135 |  |
| Tropical & Subtropical Grasslands, Savannas & Shrublands | 14854 | 7805 | 341 | 132 | 9747 | 22 |  |
| Tropical & Subtropical Moist Broadleaf Forests | 17767 | 12063 | 41673 | 1 | 45901 | 253 | 2231 |

**Table S7.** Number of VIP cells highly exposed to climate change for 2070, within each realm and biome, under scenario RCP 4.5. ‘Highly exposed’ is defined as a quarter of the Earth’s terrestrial surface with a loss of habitat condition value exceeding the 75^th^ percentile threshold for 2070, under Representative Concentration Pathway 8.5. *Very Important Places* (VIPs) span a quarter of the Earth’s terrestrial area that has the highest irreplaceability scores for tetrapods, based on 10 × 10 km grid cells, hence VIPs = 335,130 grid cells. Realms and bioregions are from Terrestrial Ecoregions of the World Version 2.0 (*85*). AT = Afrotropics, AU = Australasia, IM = Indo-Malaya, NA = Nearctic, NT = Neotropics, OC = Oceania, PA = Palearctic.

| Biomes | Biogeographic Realms |  |  |  |  |  |  |
| --- | --- | --- | --- | --- | --- | --- | --- |
|  | AT | AU | IM | NA | NT | OC | PA |
| Deserts & Xeric Shrublands | 1744 | 13 | 504 | 1122 | 525 |  | 1573 |
| Flooded Grasslands & Savannas | 4 |  | 6 |  | 163 |  |  |
| Mangroves | 441 | 298 | 669 |  | 721 |  |  |
| Mediterranean Forests, Woodlands & Scrub |  |  |  |  | 46 |  | 670 |
| Montane Grasslands & Shrublands | 1370 | 248 | 43 |  | 200 |  | 633 |
| Rock, Ice, Lakes | 1 |  |  |  | 63 |  | 151 |
| Temperate Broadleaf & Mixed Forests |  | 467 | 1286 | 315 | 1448 |  | 2842 |
| Temperate Conifer Forests |  |  | 448 | 1105 |  |  | 184 |
| Temperate Grasslands, Savannas & Shrublands | 216 |  |  | 12 | 42 |  |  |
| Tropical & Subtropical Coniferous Forests |  |  | 904 | 106 | 704 |  |  |
| Tropical & Subtropical Dry Broadleaf Forests | 218 | 85 | 1727 | 21 | 1230 | 119 |  |
| Tropical & Subtropical Grasslands, Savannas & Shrublands | 2190 | 239 | 341 | 3 | 4510 | 20 |  |
| Tropical & Subtropical Moist Broadleaf Forests | 5950 | 11180 | 29577 |  | 36520 | 240 | 226 |

**Table S8:** Percentage of Phylogenetic Diversity of terrestrial tetrapods with high exposure to climate change and/or land use impacts. *Very Important Places* (VIPs) span a quarter of the Earth’s terrestrial area that has the highest irreplaceability scores for tetrapods. Column ‘VIPs only’ represent % Phylogenetic Diversity uniquely contained within VIPs. Partitions include high exposure to climate only; high exposure to land use only; high exposure to both climate and land use; and remaining grid cells.

| Partition | RCP 4.5 |  |  |  | RCP 8.5 |  |  |  |
| --- | --- | --- | --- | --- | --- | --- | --- | --- |
|  | 2050 |  | 2070 |  | 2050 |  | 2070 |  |
|  | VIPs only | Global | VIPs only | Global | VIPs only | Global | VIPs only | Global |
| Climate only | 3.64 | 3.82 | 5.00 | 5.32 | 5.10 | 5.37 | 8.15 | 8.80 |
| Land use only | 1.78 | 1.91 | 1.29 | 1.39 | 1.51 | 1.59 | 0.25 | 0.26 |
| Climate and land use | 0.98 | 0.99 | 1.83 | 1.86 | 2.19 | 2.23 | 4.61 | 4.82 |
| Remaining | 32.87 | 93.28 | 31.15 | 91.43 | 30.47 | 90.81 | 26.27 | 86.12 |
| Total | 39.27 | 100.00 | 39.27 | 100.00 | 39.27 | 100.00 | 39.27 | 100.00 |

**Table S9:** Percentage of Species richness of terrestrial tetrapods with high exposure to climate change and/or land use impacts. *Very Important Places* (VIPs) span a quarter of the Earth’s terrestrial area that has the highest irreplaceability scores for tetrapods. Column ‘VIPs only’ represent % Phylogenetic Diversity uniquely contained within VIPs. Partitions include high exposure to climate only; high exposure to land use only; high exposure to both climate and land use; and remaining grid cells.

| Partition | RCP 4.5 |  |  |  | RCP 8.5 |  |  |  |
| --- | --- | --- | --- | --- | --- | --- | --- | --- |
|  | 2050 |  | 2070 |  | 2050 |  | 2070 |  |
|  | VIPs only | Global | VIPs only | Global | VIPs only | Global | VIPs only | Global |
| Climate only | 5.15 | 5.51 | 6.95 | 7.57 | 7.16 | 7.66 | 11.36 | 12.64 |
| Land use only | 2.52 | 2.78 | 1.77 | 1.99 | 1.97 | 2.16 | 0.29 | 0.32 |
| Climate and land use | 1.19 | 1.22 | 2.31 | 2.37 | 2.58 | 2.66 | 5.43 | 5.88 |
| Remaining | 35.69 | 90.49 | 33.53 | 88.07 | 32.83 | 87.52 | 27.47 | 81.16 |
| Total | 44.55 | 100.00 | 44.55 | 100.00 | 44.55 | 100.00 | 44.55 | 100.00 |

**Table S10:**
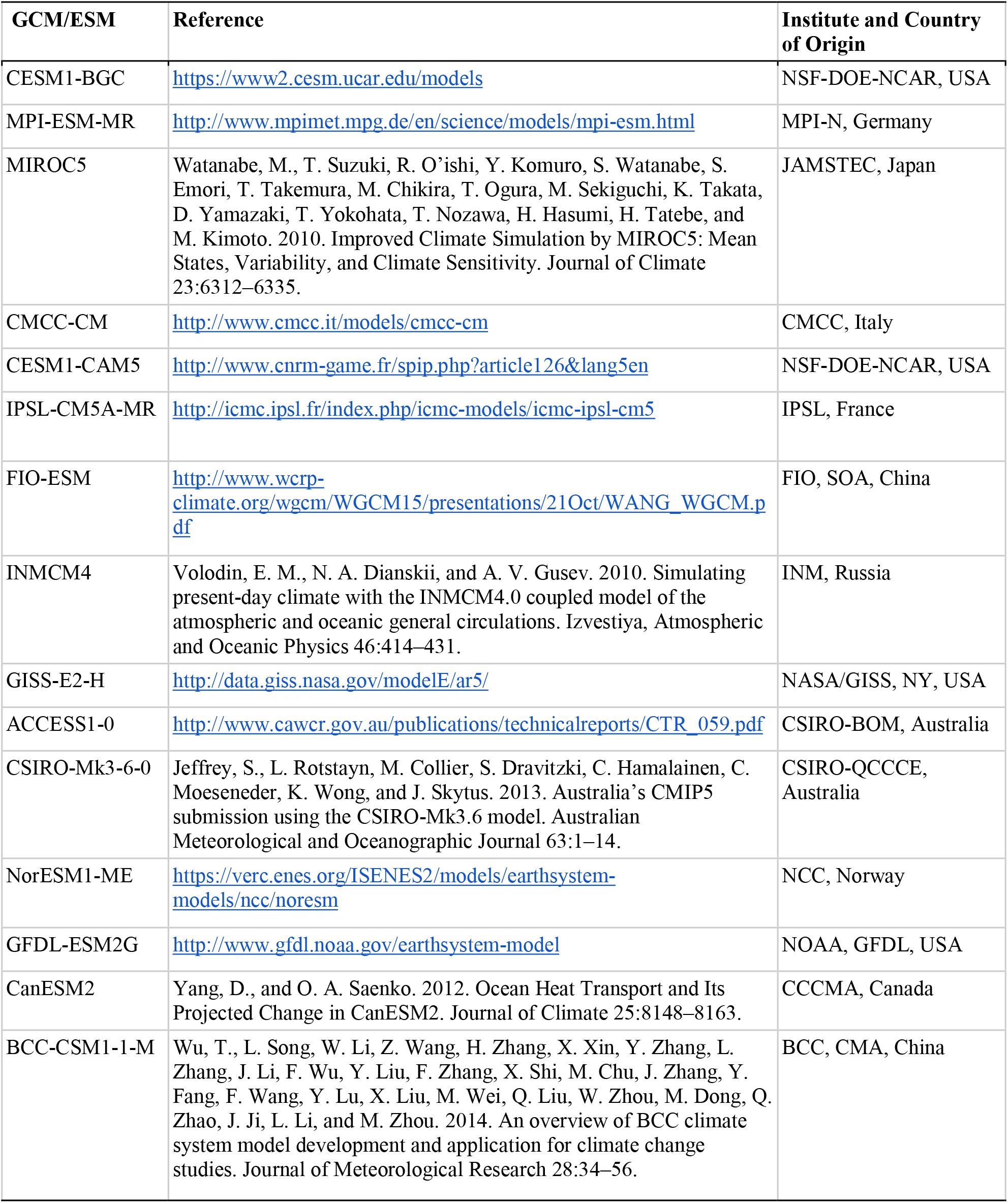
Details of the ensemble of 15 Global Climate Models (GCMs)/Earth System Models (ESMs) used to represent predicted future climate. Climate models included in this study are the highest ranked models assessed by Knutti et al. (*82*). ACCESS1-3 is Rank 3 in Sanderson et al. (*83*) but is not available from the CHELSA repository. The top sixteen models available from the CHELSA repository included the closely related ACCESS1-0 GCM, which Knutti et al. (*80*) produced very similar outputs.

| GCM/ESM | Reference | Institute and Country of Origin |
| --- | --- | --- |
| CESM1-BGC | https://www2.cesm.ucar.edu/models | NSF-DOE-NCAR, USA |
| MPI-ESM-MR | http://www.mpimet.mpg.de/en/science/models/mpi-esm.html | MPI-N, Germany |
| MIROC5 | Watanabe, M., T. Suzuki, R. O'ishi, Y. Komuro, S. Watanabe, S. Emori, T. Takemura, M. Chikira, T. Ogura, M. Sekiguchi, K. Takata, D. Yamazaki, T. Yokohata, T. Nozawa, H. Hasumi, H. Tatebe, and M. Kimoto. 2010. Improved Climate Simulation by MIROC5: Mean States, Variability, and Climate Sensitivity. *Journal of Climate* 23:6312–6335. | JAMSTEC, Japan |
| CMCC-CM | http://www.cmcc.it/models/cmcc-cm | CMCC, Italy |
| CESM1-CAM5 | http://www.cnrm-game.fr/spip.php?article126&lang5en | NSF-DOE-NCAR, USA |
| IPSL-CM5A-MR | http://icmc.ipsl.fr/index.php/icmc-models/icmc-ipsl-cm5 | IPSL, France |
| FIO-ESM | http://www.wcrp-climate.org/wgcm/WGCM15/presentations/21Oct/WANG_WGCM.pdf | FIO, SOA, China |
| INMCM4 | Volodin, E. M., N. A. Dianskii, and A. V. Gusev. 2010. Simulating present-day climate with the INMCM4.0 coupled model of the atmospheric and oceanic general circulations. *Izvestiya, Atmospheric and Oceanic Physics* 46:414–431. | INM, Russia |
| GISS-E2-H | http://data.giss.nasa.gov/modelE/ar5/ | NASA/GISS, NY, USA |
| ACCESS1-0 | http://www.cawcr.gov.au/publications/technicalreports/CTR_059.pdf | CSIRO-BOM, Australia |
| CSIRO-Mk3-6-0 | Jeffrey, S., L. Rotstayn, M. Collier, S. Dravitzki, C. Hamalainen, C. Moeseneder, K. Wong, and J. Skytus. 2013. Australia's CMIP5 submission using the CSIRO-Mk3.6 model. *Australian Meteorological and Oceanographic Journal* 63:1–14. | CSIRO-QCCCE, Australia |
| NorESM1-ME | https://verc.enes.org/ISENES2/models/earthsystem-models/ncc/noresm | NCC, Norway |
| GFDL-ESM2G | http://www.gfdl.noaa.gov/earthsystem-model | NOAA, GFDL, USA |
| CanESM2 | Yang, D., and O. A. Saenko. 2012. Ocean Heat Transport and Its Projected Change in CanESM2. *Journal of Climate* 25:8148–8163. | CCCMA, Canada |
| BCC-CSM1-1-M | Wu, T., L. Song, W. Li, Z. Wang, H. Zhang, X. Xin, Y. Zhang, L. Zhang, J. Li, F. Wu, Y. Liu, F. Zhang, X. Shi, M. Chu, J. Zhang, Y. Fang, F. Wang, Y. Lu, X. Liu, M. Wei, Q. Liu, W. Zhou, M. Dong, Q. Zhao, J. Ji, L. Li, and M. Zhou. 2014. An overview of BCC climate system model development and application for climate change studies. *Journal of Meteorological Research* 28:34–56. | BCC, CMA, China |

**Data S1. VIP-Explorer ShinyApp**. Zip file containing code and files to run Shiny-App and visualize irreplaceability of terrestrial tetrapods, and exposure to climate change and land use impacts (https://github.com/peterbat1/VIP_Explorer). The ShinyApp enables users to set different thresholds of irreplaceability and exposure to the two stressors, and explores results for all terrestrial tetrapods as well as individual clades.

**Data S2. Resolution of mismatches in species names**. The csv contains the binomial, genus, family, order, and clade names of 33,199 terrestrial tetrapods included in our phylogenetic tree and their matches to names used in species range data. Mis-matches between these datasets were identified with resolutions recorded in the csv.

**Data S3. Correlations between Complementarity of Phylogenetic Diversity, Complementarity of Species Richness, Species Richness and simple Phylogenetic Diversity**.

## Notes

### Competing Interest Statement

The authors have declared no competing interest.

